# Genomic alterations and abnormal expression of APE2 in multiple cancers

**DOI:** 10.1101/2020.01.17.910646

**Authors:** Katherine A. Jensen, Xinghua Shi, Shan Yan

## Abstract

Although APE2 plays essential roles in base excision repair and ATR-Chk1 DNA damage response (DDR) pathways, it remains unknown how the APE2 gene is altered in the human genome and whether APE2 is differentially expressed in cancer patients. Here, we report multiple-cancer analyses of APE2 genomic alterations and mRNA expression from cancer patients using available data from The Cancer Genome Atlas (TCGA). We observe that APE2 genomic alterations occur at ~17% frequency in 14 cancer types (n= 21,769). Most frequent somatic mutations of APE2 appear in uterus (2.89%) and skin (2.47%) tumor samples. Furthermore, APE2 expression is upregulated in tumor tissue compared with matched non-malignant tissue across 5 cancer types including kidney, breast, lung, liver, and uterine cancers, but not in prostate cancer. We also examine the mRNA expression of 13 other DNA repair and DDR genes from matched samples for 6 cancer types. We show that APE2 mRNA expression is positively correlated with PCNA, APE1, XRCC1, PARP1, Chk1, and Chk2 across these 6 tumor tissue types; however, groupings of other DNA repair and DDR genes are correlated with APE2 with different patterns in different cancer types. Taken together, this study demonstrates alterations and abnormal expression of APE2 from multiple cancers.

## Introduction

Genomic stability is constantly susceptible to irregularities that arise from endogenous and exogenous sources. One of the greatest threats to genomic integrity and implicated as a driving factor in numerous diseases is oxidative stress (OS), which is defined as imbalance between the production of reactive oxygen species (ROS) and antioxidant defense^1–3^. OS leads to various DNA damage such as DNA single-stranded breaks (SSBs) and DNA double-stranded breaks (DSBs). Whereas DSBs are among the most deleterious lesions, SSBs can occur upwards of 10,000 daily per cell, representing the most abundant type of DNA damage^4^. To better preserve integrity of the genome, various DNA repair and DNA damage response (DDR) pathways have evolved to mitigate the deleterious effects of OS, functioning effectively in healthy individuals^4^. It has been widely accepted that deficiencies in DNA repair and DDR pathways have been implicated in human diseases such as cancer and neurodegenerative disorders.

Previous studies have elucidated several critical proteins that regulate and coordinate various DNA repair and DDR pathways. Whereas MRN complex (Mre11, Rad50, and Nbs1) as well as BRCA1 and BRCA2 promote homology recombination (HR) -directed DSB repair^5–7^, Poly(ADP-ribose) polymerase 1 (PARP1) and X-ray repair cross-complementing protein 1 (XRCC1) have been implicated in SSB repair pathway^8^. Generally speaking, DSBs trigger the activation of ATM (ataxia-telangiectasia mutated) – Checkpoint kinase 2 (Chk2) DDR pathway^2,4^. On the other hand, SSBs and DNA replication stress trigger the activation of ATM and Rad3-related (ATR)-Checkpoint kinase 1 (Chk1) DDR pathway^9–11^. Proliferating cell nuclear antigen (PCNA) coordinates the faithful and processive DNA replication and repair of the genome when needed^12,13^. In addition to its critical role in ATR-Chk1 DDR pathway, TopBP1 participates in DSB repair directly^14–17^. Therefore, prior knowledge demonstrates that these proteins crosstalk among different DNA repair and DDR pathways.

Oxidative stress-induced DNA damage is primarily repaired by base excision repair (BER) pathway^2^. Oxidative DNA damage is first recognized and processed by various DNA glycosylases including but not limited to MUTYH, OGG1, NTH1, and NTHL1^18^. Defects in BER genes predispose to hereditary cancers. Whereas MUTYH has been established in the etiology of a colorectal cancer predisposition syndrome^19^, a recent case study reports that MUTYH germline and somatic aberrations are implicated in pancreatic ductal adenocarcinoma and breast cancer oncogenesis^20^. OGG1 is also implicated in association with lung cancer^21^. Furthermore, NTHL1 mutations or deficiencies are associated with multi-cancer phenotype including colorectal cancer and breast cancer^22^. In addition, nucleotide excision repair (NER) protein XPC may regulate OGG1 expression and participates in BER pathway to repair oxidative DNA damage, suggesting crosstalk between different DNA repair pathways^23,24^.

AP endonuclease 1 (APE1, also known as APEX1, Apn1, or Ref-1) and AP endonuclease 2 (APE2, also known as APEX2 or Apn2) have been implicated in many regulatory mechanisms in the maintenance of genome stability, especially in BER pathway^8,25,26^. In addition to its transcriptional activity of redox regulation, APE1 is the main AP endonuclease with high endonuclease activity yet weak exonuclease activity to promote BER^8,25,27^ In contrast, APE2 has high exonuclease activity but weak endonuclease activity^11,28^. APE2 is composed of three functional domains: N-terminal EEP, PCNA-interacting (PIP) motif, and a highly-conserved C-terminal zinc finger Zf-GRF^10^. APE2 interacts with PCNA via two different modes, promoting 3’-5’ SSB end resection for the activation of ATR-Chk1 DDR pathway^9–11^. Accumulating evidence has suggested that abnormal expression of APE1 is implicated in cancer such as gastric cancer and ovarian cancer^29–31^. Both APE1 and APE2 have previously been found to be upregulated in multiple myeloma cell lines^32^. However, it remains unknown whether the expression of APE2 in patient-derived tumor tissues is altered when compared with non-malignant tissues across multiple cancer types.

When genetic irregularities arise within DNA repair and DDR mechanisms, OS-induced damage may not be properly repaired, leading to genome instability and compromised protein production^2,3,33^. Copy number variations (CNVs), point mutations, and irregular gene expression patterns are examples of genomic alterations that have the potential to impact protein structure and function. Here, we conduct a multi-cancer bioinformatics analysis of APE2 at DNA, mRNA, and protein levels from publicly available data across multiple studies and cancer tissue types. In this study, we examine genomic alterations (somatic mutations and CNVs) occurring in APE2 *in vivo* from 14 cancer types and if APE2 is differentially expressed in 6 types of tumor tissue compared with non-malignant tissue. Furthermore, we analyze mRNA expression between APE2 and 13 critical DNA repair and DDR proteins. To the best of our knowledge, this work is the first to provide patient-derived evidence showing that abnormal expression of APE2 is implicated in multiple cancer types and that APE2 expression is correlated with the expression of various DNA repair and DDR proteins. We also discuss the function and biology of APE2 in genome integrity.

## Methods

### Data sets

Multiple-study data for genomic alteration events in 14 different cancer types was retrieved from the cBioPortal for Cancer Genomics. Although the cBio repository contains data for 30 different primary sites, only data for these 14 cancer types adhered to the following criteria at the time of download: (1) sample size greater than 250, (2) alteration data available and/or (3) sufficient gene expression data for the same tissue available from TCGA. Alteration events include amplifications (high-level amplification and typically focal), gains (a few additional copies but usually broad), heterozygous deletions (show loss), homozygous deletions (deep loss), and protein-level somatic mutations (docs.ciboportal.org). Study-specific details on genomic alteration event data creation can be found within individual studies. A list of all studies - with links – from which datasets have been provided is available on the web (cbioportal.org/datasets).

Gene expression quantification data of sequenced mRNA from TCGA for 6 cancer types was downloaded from Genome Data Portal (GDC) v14.0 via GDC’s transfer tool. Sample size n ≥ 20 where matched tumor and non-malignant tissue data was available for each individual was the determining factor by which cancer types were chosen for gene expression analysis. See Discussion for more information on an exception made to this criteria. Tissue samples for gene expression quantification data were originally obtained from patients at tissue source sites^34^ such as Columbia University, Mayo Clinic, Duke University, and the University of Pittsburgh. Tissue samples were then sent to one of two biospecimen collection sites (BCRs) where RNA was isolated, clinical data standardized, and analyte distribution conducted. Samples and associated data were then sent to a genome characterization or sequencing center (GCC or GSC, respectively) where data was sequenced with Illumina HiSeq technologies. For gene expression quantification, mRNA transcripts were aligned and the.bam files quantified with HTSeq. Read count data from HTSeq and from STAR are available, as well as fragments per kilobase of transcript per million mapped reads’ (FPKM) and FPKM-UQ normalized data. Read count data is often used for global differential expression analysis of gene sets. FPKM normalization method is much more specific, taking into consideration of gene length differences and is often a preferred method for gene-gene expression comparisons^35^. Since we focus on single gene and gene-gene pair analysis in this study, we use the FPKM normalized data where read count is divided by gene length.

### Data pre-processing

Genomic alteration event files containing information on APE2 per cancer type were downloaded from cBioPortal website in TSV format. Files were then programmatically parsed for relevant information from tumor tissue samples, removing duplicate data and any individuals for whom alteration event data was either not available or not explored. To reduce redundancy, only the first record for tumor tissue samples with more than one record was included in analyses.

File pre-processing for mRNA gene expression quantification included renaming files from UUID to the associated TCGA barcode. Two new sets of files were created after parsing original files: one set contained data for all tumor data for APE2 and 13 additional genes implicated in DDR pathways, one set contained only expression data from individuals for whom both tumor and matched non-malignant tissue samples were available. Table S3 shows sample sizes for tumor-only tissue and matched tissues per cancer type in our analysis.

### Genomic alteration event analysis

To characterize the frequency of genomic alteration events in APE2, the frequency of each event type per cancer type was calculated in Excel and plotted in R. Genomic alteration plots were created in R and figures representing APE2 protein domains and mutations – along with R-generated plots – were created in PowerPoint.

### mRNA analysis

To find the difference in mRNA expression values of APE2 between tumor and matched non-malignant tissue, a paired two-sided t-test was conducted and boxplots were created in R. To find correlations between mRNA expression of APE2 to other genes implicated in DDR pathways, FPKM values were log-transformed, Pearson’s correlation run and scatterplots created in R.

### Bioinformatics tools

Data from the GDC portal was downloaded using the provided transfer tool. For pre-processing of data, scripts written in Python, utilizing the *os.walk* module, were used to parse, create, and rename files. To find matching TCGA barcode per UUID, *UUIDtoBarcode* from the R-package TCGAUtils was used^36^. From R-package *ggplot2* and dependencies, genomic alteration frequency plot and somatic mutation plots were created using *ggplot* and scatterplots created with *ggscatter*^37^. All R-packages were implemented, and statistical analyses and plots were created, with RMarkdown in RStudio 1.1.456^37–41^. All Python scripts were run on Ubuntu 18.04LTS from Windows 64-bit OS. Figures created in PowerPoint were done so using Microsoft Office 2019 PowerPoint Version 1912 (Build 12325.20240 Click-to-Run).

## Results

### Genomic alterations in APE2 across all cancer types

To verify the occurrence of APE2-situated genomic alterations in cancer patients, data from 14 cancer-types in the cBiolPortal database was programmatically analyzed. From all samples for which data were available and had been evaluated for genomic alterations (n=21,769), genomic alterations in APE2 occurred at ~17% frequency and appeared in each cancer type (Fig. 1). Cancers with the highest frequency of total events were skin (24.35%), liver (23.88%), and breast (23.72%) (Fig. 1, Table S1). From the total alteration events observed, heterozygous deletions occurred most frequently (51.18%), gains next (40.77%), then amplifications (3.77%) and mutations (3.11%) with homozygous deletions occurring at the lowest frequency (1.17%). CNVs in APE2 were found at varying levels across each cancer type. Gains occurred most frequently in lymphoid cancer (14.30%), amplifications most often in prostate cancer (3.28%), heterozygous deletions observed most frequently in liver tumor tissue (15.42%), and head and neck cancerous samples revealed the highest frequency of homozygous deletions (1.07%) (Fig. 1, Table S1). Frequency of somatic mutations in APE2 ranged from 2.89% (uterus) to 0.10% (head and neck), although its somatic mutations did not appear in kidney nor pancreas tumor tissue.

**Figure 1.**
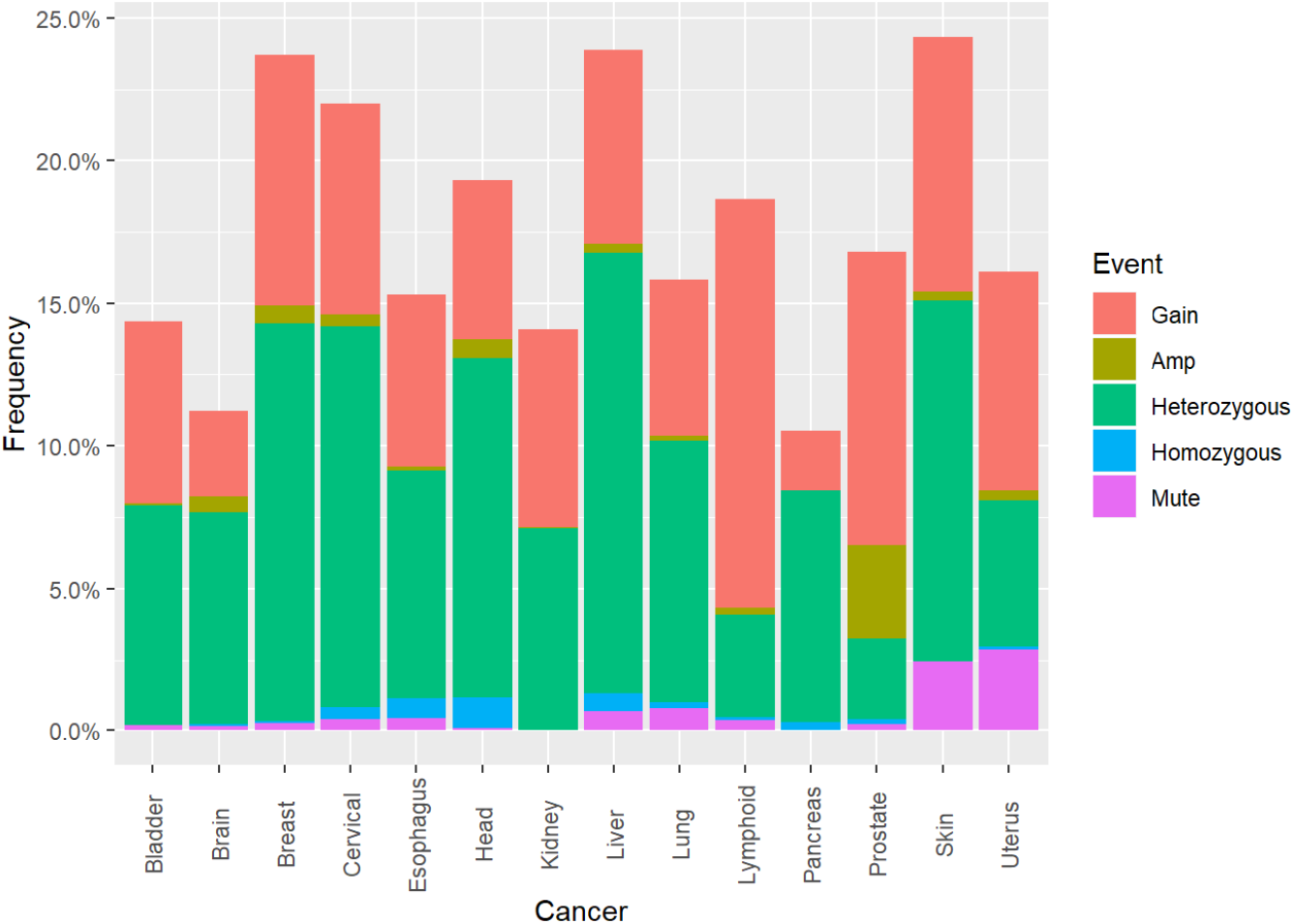
Frequency of genomic alteration events in APE2 across 14 cancer types and its mRNA expression analysis between tumor tissue and matched non-malignant tissue from 6 different cancer types. cBioPortal analysis of APE2 genomic alterations including gains (Gain), amplifications (Amp), heterozygous deletions, homozygous deletions, and somatic mutations (Mute). Redundant samples were removed before analysis. Multiple somatic mutation events within a single sample and duplicate mutations across different individuals were each distinctly counted.

### Occurrences of coding truncations and amino acid changes in APE2

In addition to the analysis of APE2 genomic alterations, protein consequences of somatic mutations in APE2 were analyzed. Protein-level annotations of somatic mutations in APE2 were extracted from genomic alteration event data for 12 cancer types (Table S2). A total of 117 mutations were found in APE2 with uterine (40), lung (22), and skin (16) tumor tissue revealing the highest number of mutations (Fig. 2A, Table S2). Mutation events were comprised mainly of missense mutations with several nonsense mutations which create premature stop codons, frameshift deletions which are deletions of chunks of protein sequence that create a shift in remaining amino acids, and alternative splices denoted with an ‘X’ indicating a translation termination codon at the site (Fig. 2A–2C). Tumor tissues with the highest frequency of somatic mutations in APE2 were uterus (2.89%= 40 out of 1,386), skin (2.47%= 16 out of 649) and lung (0.78%= 22 out of 2,831) tumor samples (Fig. 2B). Notably, out of the 117 mutations, 29 Arginine residues (~25%) were mutated to other residues, suggesting a distinct feature of APE2 missense mutations (Fig. 2A–2B, Table S2). The Arginine residue at position 465 was mutated to Histidine (i.e., R465H) in skin and cervical tumor samples, and to Cysteine (i.e., R465C) in more than one individual with uterine cancer. An Arginine to Cysteine mutation was found at position 173 (i.e., R173C) in one breast cancer and two uterine tumor samples. A R222C and a R222H mutations were found in uterine cancer, and the R222C mutation was also found in a brain cancer sample.

**Figure 2.**
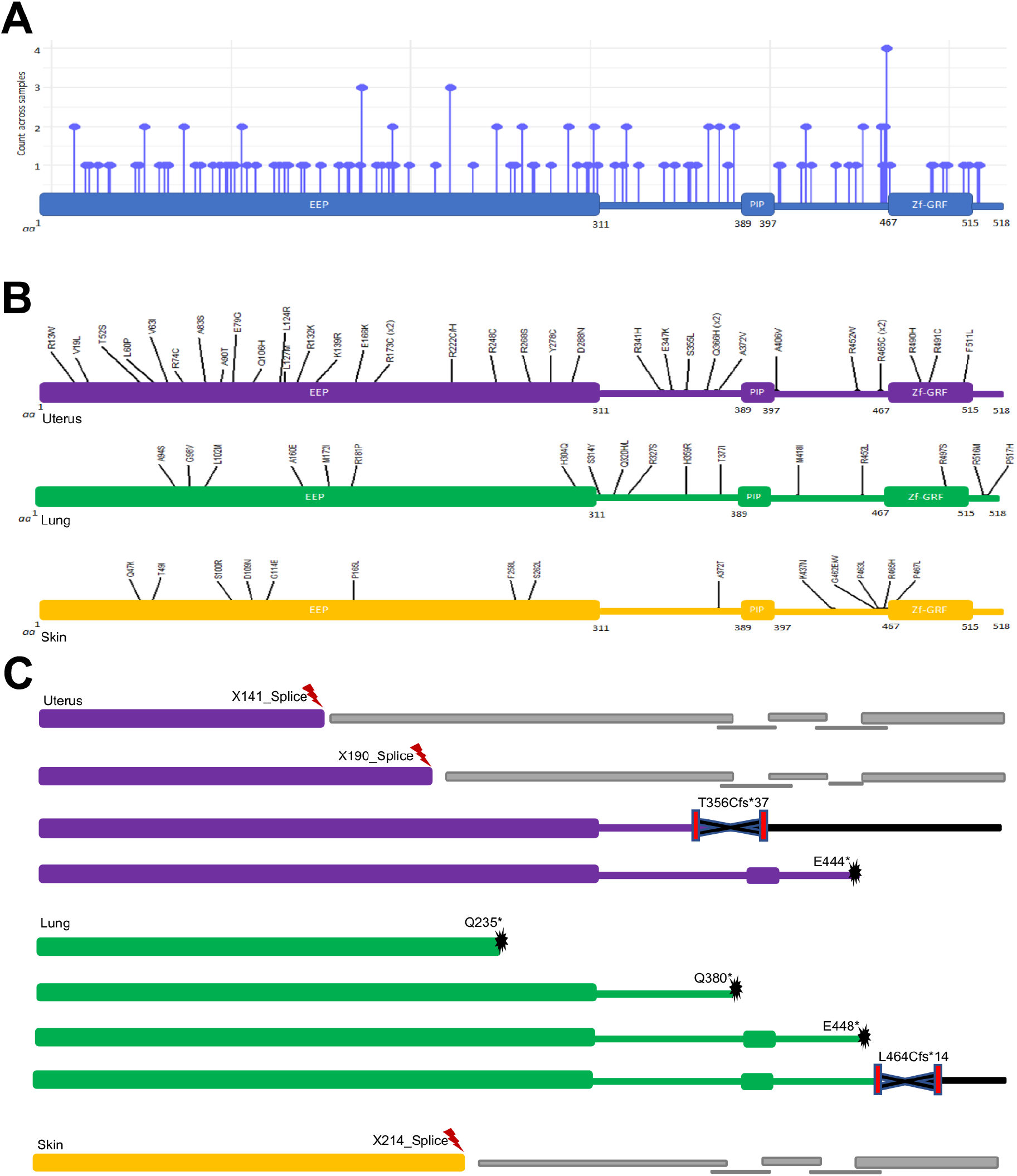
Somatic mutations in APE2 at the protein level across multiple cancer types. (**A**) mutation counts of different mutations in APE2 protein in 12 different cancer types. (**B**) missense mutations in APE2 protein in uterus, skin, and lung cancers. (**C**) representative splicing and truncations in APE2 protein in uterus, lung, and skin cancers. The images were created by *R* and PowerPoint software (see Methods section for details).

Additionally, four nonsense mutations of APE2 were found in lung cancers (Q235*, Q380*, and E448*) and uterine cancer (E444*) creating premature stop codons (Fig. 2C). Of the two frameshift deletions found, one was in uterine tissue (T356Cfs*37) and one in lung cancer (L464Cfs*14) (Fig. 2C). Furthermore, three alternative splices were revealed in uterus (X141, X190) and skin cancers (X214) (Fig. 2C).

### Upregulation of APE2 mRNA expression in tumor tissue

After establishing the occurrence of alterations in APE2 at the genomic level, APE2 expression at the mRNA level was analyzed. To find if APE2 is differentially expressed in cancer patients, gene expression quantification data was computationally compared between tumor and non-malignant tissue using a two-sided *t*-test in *R*. Samples referred to as ‘matched’ indicate individuals where non-malignant and tumor tissue were both available in the data (Table S3 and Table S4). FPKM (fragments per kilobase of exon model per million reads mapped) values of mRNA sequencing data for APE2 in matched samples revealed significant upregulation (*α* = .*05*) in tumor tissue compared with non-malignant tissues across 5 cancer types including kidney (n = 126), breast (n = 112), lung (n = 106), liver (n = 58), and uterine (n = 23) cancers (Fig. 3A–3D, 3F). Matched samples from prostate cancer patients (n = 52) did not present a significant difference in APE2 FPKM values (Fig. 3E).

**Figure 3.**
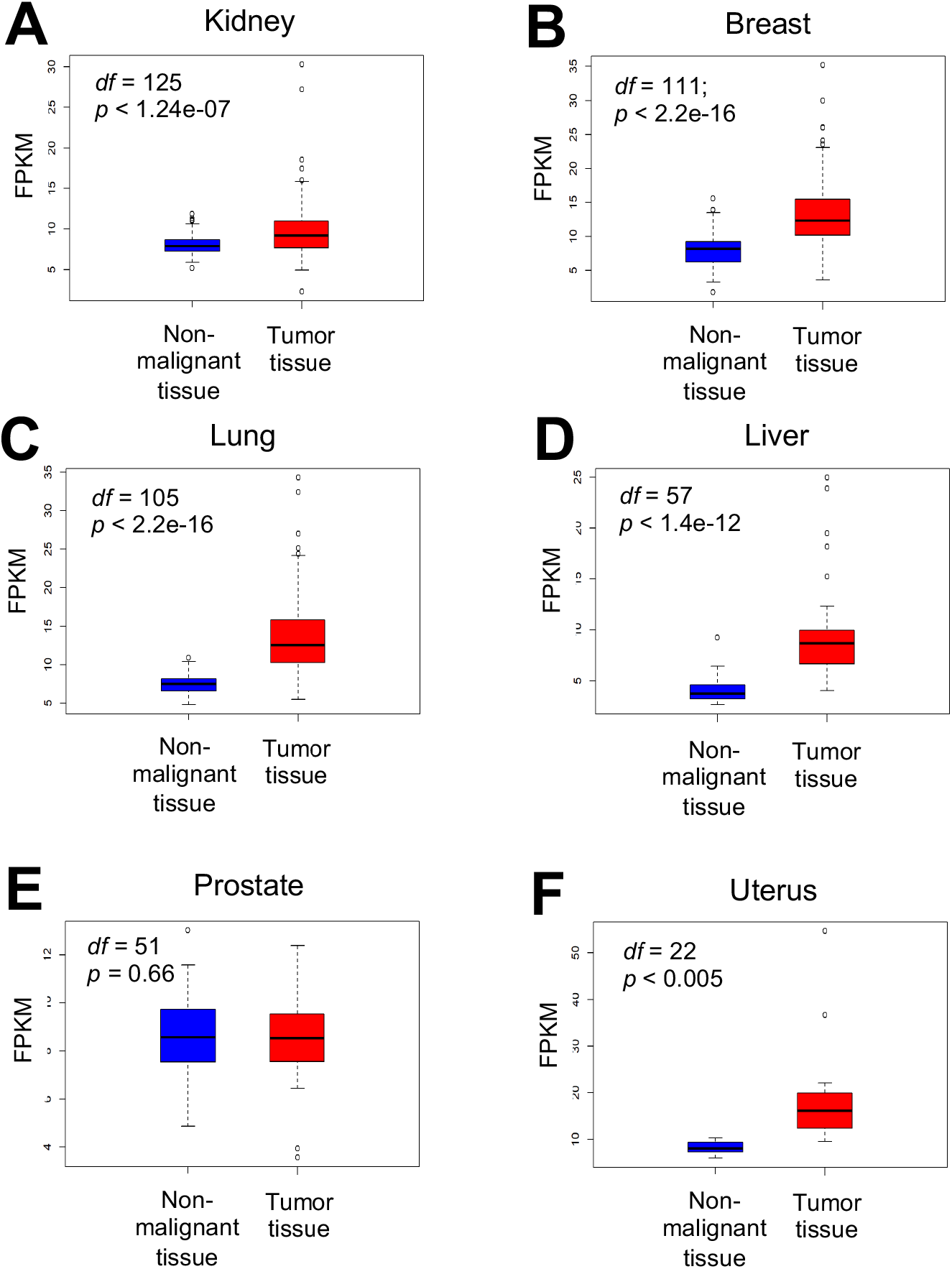
APE2 mRNA expression between tumor tissue and matched non-malignant tissue per individual from 6 different cancer types including kidney (**A**), breast (**B**), lung (**C**), liver (**D**), prostate (**E**), and uterus (**F**). A paired two-sided *t*-test was conducted to determine significance of difference. *df* (degree of freedom) and *p* values are listed in each panel.

To establish a baseline for finding correlations - between expression patterns in APE2 and 13 other DNA repair and DDR pathway genes – in tumor tissue, matched samples were further analyzed for differential expression in these genes. Notably, in most, if not all, cancer types analyzed, mRNA expression of 11 out of the 13 DNA repair and DDR genes are upregulated in tumor tissue compared with that in non-malignant tissue. Analysis of these genes revealed significant upregulation in APE1, BRCA1, Chk1, Chk2, and TopBP1 for all cancer types (Fig. S1-S5). Like APE2, PCNA, BRCA2 and XRCC1 were also significantly upregulated in all cancer types except prostate (Fig. S6-S8). PARP1 was upregulated in all cancer types except kidney cancer (Fig. S9). mRNA expression of ATR in tumor tissue was increased compared with matched non-malignant tissue in liver, lung, uterus, and prostate cancers, but not in breast and kidney cancers (Fig. S10). Rad50 mRNA expression was upregulated in almost all cancer types except uterine cancer (Fig. S11).

Intriguingly, mRNA expression of ATM and Mre11 was upregulated in tumor tissue compared with matched samples in some cancer types but downregulated in other cancer types (Fig. S12-S13). ATM expression was upregulated in kidney and liver cancers, but not in lung and prostate cancers, while it was downregulated in breast and uterine cancers (Fig. S12). Mre11 expression was upregulated in lung and liver cancers, but not in kidney, prostate, and uterus cancers, and was significantly downregulated in breast cancer (Fig. S13).

### Patterns in mRNA expression between APE2 and other DNA repair and DDR genes

Because co-expression and interactive partners of APE2 are not yet well known, it is significant to compare expression levels of APE2 to other DNA repair and DDR genes. After establishing differential expression patterns for reach gene per cancer types, FPKM values of APE2 were correlated with 13 other DNA repair and DDR genes in tumor tissue only for breast (n= 1,105), lung (n= 1,028), kidney (n= 891), uterine (n= 552), prostate (n= 499), and liver (n= 407) cancers (Table S3 and S5). While APE2 had a significant (α= 0.05) positive correlation with PCNA, APE1, XRCC1, PARP1, Chk1, and Chk2 across all 6 cancer types, groupings of DNA repair and DDR genes were found to be correlated with APE2 in different patterns in different cancer types (Fig. 4, and Fig. S14-S18). In liver cancer, correlations of APE2 with these 13 DNA repair and DDR genes were all positive (Fig. S14). Out of the 13 other genes, 11 had the significantly strongest relationship per gene with APE2 in liver cancer (determined by *R*-value and α= 0.05) (Fig. S14); however, direction, strength, and significance of correlations varied in other cancer types.

**Figure 4.**
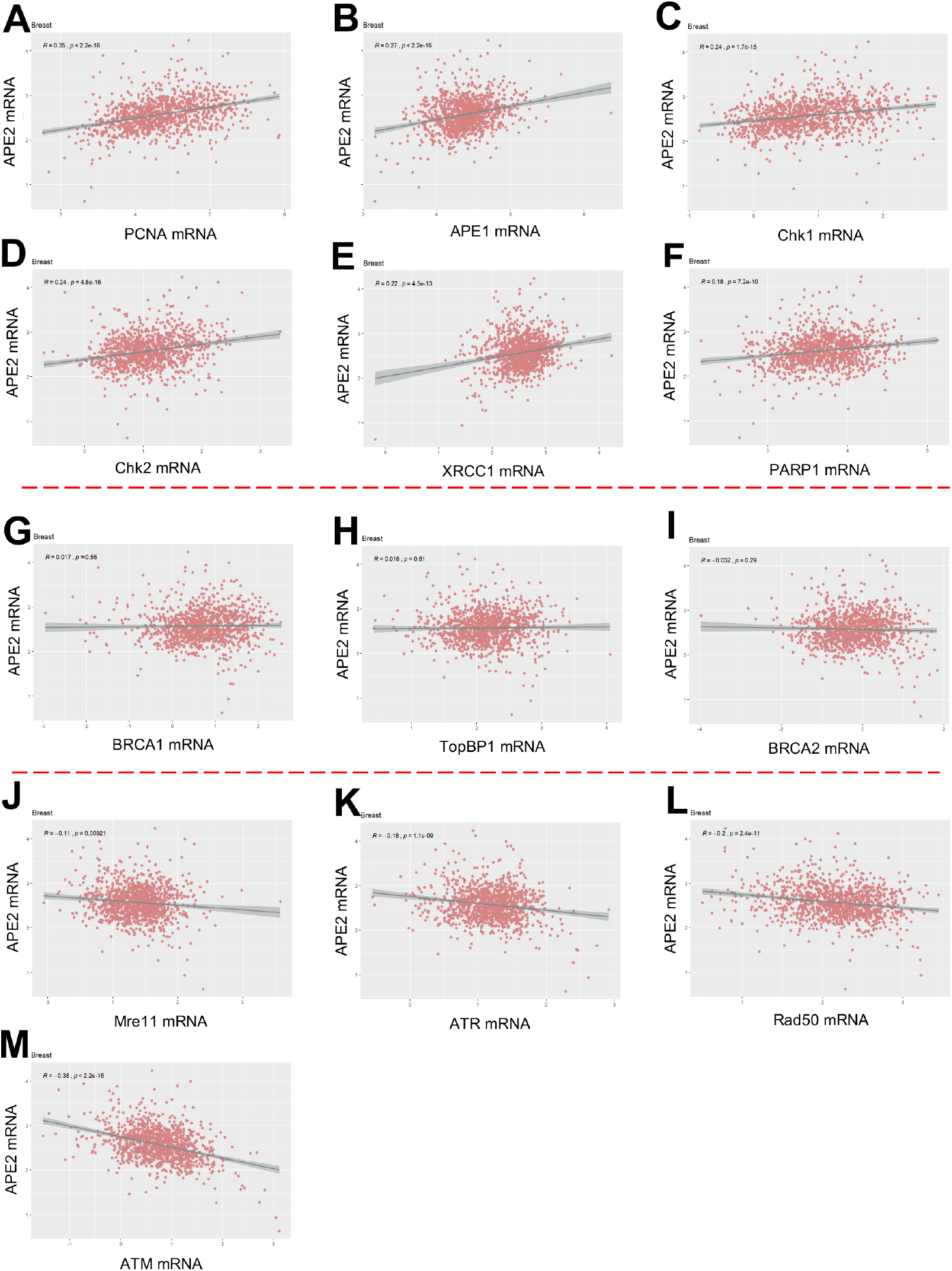
Correlation between mRNA expression of APE2 and 13 other DNA repair and DDR proteins in tumor tissues of breast cancer. (**A-F**) are positive correlation of APE2 mRNA expression with that of PCNA (**A**), APE1 (**B**) Chk1 (**C**), Chk2 (**D**), XRCC1 (**E**), and PARP1 (**F**). (**G-I**) No correlation of APE2 mRNA expression with that of BRCA1 (**G**), TopBP1 (**H**), and BRCA2 (**I**). (**J-M**) show negative correlation of APE2 mRNA expression with that of Mre11 (**J**), ATR (**K**), Rad50 (**L**), and ATM (**M**). Pearson’s *R* and *p* values are listed in each panel.

#### PCNA, APE1, XRCC1, and PARP1

Across all 6 cancer types, APE2 expression was significantly positively correlated with the expression of PCNA, APE1, PARP1 and XRCC1 yet expression ranges varied per cancer type and gene-gene pair. The positive correlation between the expression of APE2 and PCNA was most significant in prostate cancer. However, neither APE2 nor PCNA expression values was significantly different in matched prostate tumor (Fig. 3E and Fig. S6E), thus there is no baseline to strengthen the implications of this APE2-PCNA correlation. Weak APE2-APE1 expression correlations were observed in lung and kidney cancers, which is consistent with the significant role of APE1 and APE2 in SSB repair^11,42,43^. When APE2 and PARP1 values were correlated, liver and uterine cancer samples showed more variability whereas very little spread occurred in breast and kidney cancer samples. The highest positive correlation of APE2 and XRCC1 occurred in liver cancer.

#### Chk1 and Chk2

APE2 mRNA expression was positively and significantly correlated with the mRNA expression of Chk1 and Chk2 in all 6 cancer types (breast, liver, prostate, uterine, lung, and kidney) (Fig. 4, Fig. S14-S18). It is noted that the strongest positive correlation of APE2 with Chk1 and Chk2 is found in lung cancer (Fig. S17).

#### BRCA1 and BRCA2

When APE2 expression was correlated with expression of BRCA1 and BRCA2, the top 4 strongest positive correlations were, in order starting with strongest, liver, lung, prostate, and uterus. Whereas there were no significant correlations of APE2 to BRCA1 and BRCA2 in breast cancer (Fig. 4), kidney cancer samples exhibited a negative correlation of APE2-BRCA2 and a positive correlation of APE2-BRCA1 (Fig. S18, and Table S5).

#### TopBP1 and ATR

The APE2-ATR relationship was positive in liver and lung cancer but negative in breast cancer. Furthermore, the APE2-ATR correlation was not significant in prostate, uterine, nor kidney cancers. Although a positive correlation of APE2-TopBP1 was revealed in liver, prostate, uterus, and lung cancers, the APE2-TopBP1 correlation was not significant in kidney nor breast cancer.

#### ATM, Mre11, and Rad50

The APE2-ATM correlation was negative in all cancer types expect liver. APE2 did not have a significant relationship with Mre11 or Rad50 in uterine nor prostate cancers. The correlation of APE2 with Mre11 was positive in liver and lung cancers, and negative in breast and kidney cancers. In addition, the APE2-Rad50 correlation was negative in breast, kidney, and lung cancers, yet positive in liver cancer.

## Discussion

### Function and biology of APE2

The human APE2 gene was first cloned and briefly characterized in 2000^44^. However, the understanding of APE2 in DNA repair and DDR pathways has been derived from studies of model organisms such as *Xenopus* and yeast^9–11,45–49^, despite some biochemical and sub-cellular localization characterization of human APE2 protein^28,50,51^. Recent genetic screens identified APE2 as a synthetic lethal target in BRCA1- and BRCA2-deficient colonic and ovarian cancer cell lines^52^. Although the exact underlying mechanism remains unknown, this report suggests that APE2 may contribute to different DNA repair pathways other than BRCA1- and BRCA2-mediated DSB repair. Consistent with this, our series of studies using *Xenopus* egg extract system suggest that APE2 plays a direct role in SSB repair via the 3’-5’ SSB end resection^11,26,43^. Of note, gene expression of APE2 and APE1 has been found up-regulated in multiple myeloma (MM) patients and MM cells, which may lead to dysregulation of HR via regulating Rad51 expression^32^. These new evidences suggest that APE2 contributes to genome integrity via different mechanisms.

A prior study has shown that APE2-knock out (KO) mice are viable but develop immune response defects and growth retardation^53^. Subsequent characterization of APE2-KO mice revealed the significance of APE2 in B cell development and immunoglobulin class switch recombination^54,55^. Interestingly, recent evidences suggest that ATR and BER pathways are involved in regulating the expression of Programmed death-ligand 1 (PD-L1), an important player for immunotherapy^56–58^. As APE2 has been shown in both BER and ATR pathways^11,50^, it is interesting to test whether APE2 is directly involved in cancer immunotherapy in future studies.

Due to the lack of in-dept knowledge about APE2 functions in human diseases, the purpose of this study has been to contribute knowledge from such an angle to investigate the genomic alterations and abnormal expression of APE2 in human cancers. Using data from samples across multiple cancer types has been advantageous in providing a snapshot of APE2 characteristics in cancer yet there are limitations which must also be acknowledged.

## Genomic alterations and abnormal expression of APE2 in cancer samples

Our data from 21,769 cancer patients revealed that genomic alterations of APE2 occur with approximately 17% frequency across all cancer types (10.54% in pancreas - 24.35% in skin). This observation suggests the potential involvement of APE2 in cancer development, although future studies are needed to test this question directly. Frequencies of CNVs in APE2 for gains, amplifications, heterozygous deletions, and homozygous deletions vary significantly per cancer type. For example, the frequencies of APE2 gain (8.94%), amplifications (0.31%), heterozygous deletions (12.63%), and homozygous deletions (0%) from 1,386 skin cancer patients are different from those frequencies in other cancer types. In some cancers, such as breast and liver, heterozygous deletions are the most frequent CNV type. Alternatively, for lymphoid, uterine, and prostate cancer, gains occur most frequently.

In addition, Arginine mutations dominating APE2 somatic mutations are a distinct feature. In particular, there are 8 Arginine mutations localized in the extreme C-terminus Zf-GRF motif of APE2 (i.e., R465C, R465H, R490H, R491C, R497S, R499W, R508Q, R516M). Our previous studies have shown that R473A, R473E, R502A, and R502E mutant APE2 in *Xenopus* (homologous to R479 and R508 mutants in human APE2) is deficient in ssDNA binding^10^, and that G483A-R484A double mutants of *Xenopus* APE2 (homologous to G489A-R490A double mutants in human APE2) are defective for ssDNA binding and PCNA interaction^11^. We speculate that the Arginine missense mutations in human APE2 identified from this study may compromise its ssDNA binding and/or PCNA interaction, leading to compromised exonuclease activity and associated genome instability.

Our results on APE2 mRNA expression from matched tumor and non-malignant tissue demonstrate its overexpression in kidney, breast, lung, liver, and uterine cancers, but not in prostate cancer (Fig. 3). This pattern of APE2 overexpression is also found in PCNA, BRCA2, and XRCC1 (Fig. S6-S8). Consistent with our findings, Kumar et al. recently reported that APE2 is overexpressed at the levels of mRNA and protein in multiple myeloma (MM) cell lines and 112 MM patients from two datasets^32^. Previous studies have demonstrated that PCNA interacts with APE2 to regulate its exonuclease activity^11,28,45,47,50^. A recent genetic screen has revealed that APE2 is a synthetic lethal target in BRCA2-deficient cells^52^. Although APE2 and XRCC1 are involved in BER pathway, it seems that they contribute to SSB repair via different mechanisms^43^.

A prior study revealed that mRNA expression of APE1 and PARP1 is upregulated in tumor tissue compared with that in non-malignant tissues from 53 paired colorectal cancer patients^59^. This is consistent with our observation of upregulation of APE1 mRNA in kidney, breast, lung, liver, prostate and uterus cancers (Fig. S1), and increased PARP1 mRNA expression in breast, lung, liver, prostate and uterus cancers (Fig. S9). Although upregulation or downregulation of individual genes in the BER pathway is often found in tumor tissue over non-malignant tissue, overall BER capacity has been proposed a determinant of prognosis and therapy response to DNA-damaging 5-fluorouracial in colon cancer patients^60^.

## Correlation of APE2 mRNA expression with other DNA repair and DDR proteins

We have shown that APE2 expression is positively correlated with the expression of PCNA, APE1, PARP1, XRCC1, Chk1, and Chk2 across all six cancer types analyzed in this work (Fig. 4, Fig. S14-S18). This may be interpreted that APE2 is involved in BER and SSB repair pathways as same as PCNA, APE1, XRCC1, and PARP1^11,28,42,43^. Recent studies showing the role of APE2 in Chk1 phosphorylation in oxidative stress could provide feasible explanation for the positive correlation found between APE2 and Chk1^9,10^. Interestingly, APE2 expression is negatively correlated with ATM expression in all cancer types except liver cancer, suggesting that APE2 and ATM may be in different DDR pathways.

Due to tumor heterogeneity, there is no ‘one-size-fits-all’ formula that can be applied to the prediction, prevention, or therapy of cancer or other human diseases. In this study we demonstrate the varying gene expression and genomic alteration profiles of APE2 by tissue site and even by individual in multiple cancers. Future studies that include more tissue sites, additional metadata, and consider other potential molecular partners of APE2 will be useful in establishing the genomic landscape in which certain APE2 characteristics can be implicated as predictive markers or therapeutic targets.

## Consideration of sample sizes

The presence of matched samples in TCGA dataset provides clear indication of whether APE2 changes in tumor tissue per individual, yet many of the cancer type sample sizes of matched tissues in TCGA were too small to provide statistical value. Due to the prevalence of some cancer types over others, it is infeasible to obtain consistently large quantities of data for all cancer types. Furthermore, some tumor tissue samples are available in ‘study-worthy’ amounts, yet the collection of matched non-malignant tissues is not available due to the nature of the tissue. For instance, brain cancer datasets contain little to no matched non-malignant samples because extraction of non-malignant tissue from a living human brain could have detrimental effects on the individual. Additionally, tumor location and size can impact the availability of non-malignant tissue. If the affected organ is already small or the tumor has spread to the entire organ, insufficient amounts of non-malignant tissue would be left for extraction or analysis.

## Consideration of data availability

Gene expression quantification data was useful in understanding the transcriptome surrounding APE2. In this study, data from multiple cancer types reveals not only APE2 characteristics in disease but suggests tissue-dependent or tissue-specific characteristics. All available data from TCGA has been subject to uniform protocol and strict quality control, indications of valid and consistent datasets. However, data from certain cancer studies could not be used in this version of TCGA data due to discrepancies. For example, bladder and colon cancer datasets contained identical gene expression quantification data and thus, due to lack of clarity, were not included in the current study. Protein expression data from The Cancer Proteome Atlas (TCPA) is available for many of the same individuals in the TCGA datasets we have analyzed here. An analysis of matched protein expression could be duly advantageous for examining APE2 transcription levels to the level of expressed protein and to compare translation of APE2 with other DNA repair and DDR genes. Regardless, APE2 is not included in the TCGA/TCPA protein quantification analyses, deeming the protein expression data inefficient for the current study.

Downloadable files of somatic mutation data from cBio contained conflicting information from that which could be visualized on the web interface of cBioPortal. For example, somatic mutations A372T and G462E found in skin cancer were not included in the files but do appear on the cBioPortal website. These missing mutations were found and manually added to analyses. Likewise, genomic alteration data from cBio is missing for a large number of samples, resulting in slightly less robust results. Furthermore, ploidy and purity of copy number calls could be different across studies. Due to a general lack of prior manual review that could ensure uniform quality of data available from cBioPortal, the CNV data has the potential to represent a number of false positives and false negatives (docs.cbiopotal.org). Despite minor limitations with sample sizes and lack of availability for particular data-types, we believe the magnitude and overall quality of data analyzed here has been sufficient to capture certain patient-derived characteristics of APE2 in disease.

## Acknowledgements

We thank the members from the Yan lab and the Shi lab for critical comments during early stage of the study. The Yan lab is supported, in part, by funds from UNC Charlotte and grants from the NIH/NCI (R01CA225637) and NIH/NIGMS (R15GM114713). The Shi lab is supported by NIH/NHGRI (R15HG009565). K.J. was supported by the Charlotte Research Scholar Program in University of North Carolina at Charlotte.

## Authors contributions

K.J., X.S., and S.Y. designed experiments. K.J. performed experiments. K.J., X.S., and S.Y. analyzed the data. K.J., X.S., and S.Y. wrote the manuscript.

## Availability of data and materials

The gene expression quantification datasets analyzed during the current study are publicly available at https://portal.gdc.cancer.gov/repository. Genomic alteration data is publicly available at https://www.cbioportal.org/. GDC’s transfer tool can be downloaded from https://gdc.cancer.gov/access-data/gdc-data-transfer-tool. Code is available at https://github.com/shilab/cancer-data.

## Additional information

Correspondence and request for materials should be addressed to S.Y. and X.S.

## References

1 Hoch, N. C. et al. XRCC1 mutation is associated with PARP1 hyperactivation and cerebellar ataxia. Nature 541, 87–91, doi:10.1038/nature20790 (2017).

2 Yan, S., Sorrell, M. & Berman, Z. Functional interplay between ATM/ATR-mediated DNA damage response and DNA repair pathways in oxidative stress. Cell Mol Life Sci 71, 3951–3967, doi:10.1007/s00018-014-1666-4 (2014).

3 Betteridge, D. J. What is oxidative stress? Metabolism 49, 3–8, doi:10.1016/s0026-0495(00)80077-3 (2000).

4 Caldecott, K. W. Single-strand break repair and genetic disease. Nat Rev Genet 9, 619–631, doi:10.1038/nrg2380 (2008).

5 Chen, C. C., Feng, W., Lim, P. X., Kass, E. M. & Jasin, M. Homology-Directed Repair and the Role of BRCA1, BRCA2, and Related Proteins in Genome Integrity and Cancer. Annu Rev Cancer Biol 2, 313–336, doi:10.1146/annurev-cancerbio-030617-050502 (2018).

6 Moynahan, M. E., Pierce, A. J. & Jasin, M. BRCA2 is required for homology-directed repair of chromosomal breaks. Mol Cell 7, 263–272, doi:10.1016/s1097-2765(01)00174-5 (2001).

7 Zhao, W. et al. BRCA1-BARD1 promotes RAD51-mediated homologous DNA pairing. Nature 550, 360–365, doi:10.1038/nature24060 (2017).

8 Abbotts, R. & Wilson, D. M., 3rd. Coordination of DNA single strand break repair. Free Radic Biol Med 107, 228–244, doi:10.1016/j.freeradbiomed.2016.11.039 (2017).

9 Willis, J., Patel, Y., Lentz, B. L. & Yan, S. APE2 is required for ATR-Chk1 checkpoint activation in response to oxidative stress. Proc Natl Acad Sci USA 110, 10592–10597, doi:10.1073/pnas.1301445110 (2013).

10 Wallace, B. D. et al. APE2 Zf-GRF facilitates 3’-5’ resection of DNA damage following oxidative stress. Proc Natl Acad Sci USA 114, 304–309, doi:10.1073/pnas.1610011114 (2017).

11 Lin, Y. et al. APE2 promotes DNA damage response pathway from a single-strand break. Nucleic Acids Res 46, 2479–2494, doi:10.1093/nar/gky020 (2018).

12 Choe, K. N. & Moldovan, G. L. Forging Ahead through Darkness: PCNA, Still the Principal Conductor at the Replication Fork. Mol Cell 65, 380–392, doi:10.1016/j.molcel.2016.12.020 (2017).

13 Slade, D. Maneuvers on PCNA Rings during DNA Replication and Repair. Genes (Basel) 9, doi:10.3390/genes9080416 (2018).

14 Liu, Y. et al. TOPBP1(Dpb11) plays a conserved role in homologous recombination DNA repair through the coordinated recruitment of 53BP1(Rad9). J Cell Biol 216, 623–639, doi:10.1083/jcb.201607031 (2017).

15 Ogiwara, H. et al. Dpb11, the budding yeast homolog of TopBP1, functions with the checkpoint clamp in recombination repair. Nucleic Acids Res 34, 3389–3398, doi:10.1093/nar/gkl411 (2006).

16 Kumagai, A., Lee, J., Yoo, H. Y. & Dunphy, W. G. TopBP1 activates the ATR-ATRIP complex. Cell 124, 943–955, doi:10.1016/j.cell.2005.12.041 (2006).

17 Yan, S. & Michael, W. M. TopBP1 and DNA polymerase alpha-mediated recruitment of the 9-1-1 complex to stalled replication forks: implications for a replication restart-based mechanism for ATR checkpoint activation. Cell Cycle 8, 2877–2884, doi:10.4161/cc.8.18.9485 (2009).

18 Krokan, H. E. & Bjoras, M. Base excision repair. Cold Spring Harb Perspect Biol 5, a012583, doi:10.1101/cshperspect.a012583 (2013).

19 Banda, D. M., Nunez, N. N., Burnside, M. A., Bradshaw, K. M. & David, S. S. Repair of 8-oxoG:A mismatches by the MUTYH glycosylase: Mechanism, metals and medicine. Free Radic Biol Med 107, 202–215, doi:10.1016/j.freeradbiomed.2017.01.008 (2017).

20 Thibodeau, M. L. et al. Base excision repair deficiency signatures implicate germline and somatic MUTYH aberrations in pancreatic ductal adenocarcinoma and breast cancer oncogenesis. Cold Spring Harb Mol Case Stud 5, doi:10.1101/mcs.a003681 (2019).

21 Vlahopoulos, S., Adamaki, M., Khoury, N., Zoumpourlis, V. & Boldogh, I. Roles of DNA repair enzyme OGG1 in innate immunity and its significance for lung cancer. Pharmaco Ther 194, 59–72, doi:10.1016/j.pharmthera.2018.09.004 (2019).

22 Grolleman, J. E. et al. Mutational Signature Analysis Reveals NTHL1 Deficiency to Cause a Multi-tumor Phenotype. Cancer Cell 35, 256–266 e255, doi:10.1016/j.ccell.2018.12.011 (2019).

23 de Melo, J. T. et al. XPC deficiency is related to APE1 and OGG1 expression and function. Mutat Res 784-785, 25–33, doi:10.1016/j.mrfmmm.2016.01.004 (2016).

24 Melis, J. P. et al. Slow accumulation of mutations in Xpc(-/-) mice upon induction of oxidative stress. DNA Repair (Amst) 12, 1081–1086, doi:10.1016/j.dnarep.2013.08.019 (2013).

25 Tell, G., Quadrifoglio, F., Tiribelli, C. & Kelley, M. R. The many functions of APE1/Ref-1: not only a DNA repair enzyme. Antioxid Redox Signal 11, 601–620, doi:10.1089/ars.2008.2194 (2009).

26 Hossain, M. A., Lin, Y. & Yan, S. Single-Strand Break End Resection in Genome Integrity: Mechanism and Regulation by APE2. Inter J Mol Sci 19, 2389, doi:10.3390/ijms19082389 (2018).

27 Wilson, D. M., 3rd. Properties of and substrate determinants for the exonuclease activity of human apurinic endonuclease Ape1. J Mol Biol 330, 1027–1037, doi:10.1016/s0022-2836(03)00712-5 (2003).

28 Burkovics, P., Hajdu, I., Szukacsov, V., Unk, I. & Haracska, L. Role of PCNA-dependent stimulation of 3’-phosphodiesterase and 3’-5’ exonuclease activities of human Ape2 in repair of oxidative DNA damage. Nucleic Acids Res 37, 4247–4255, doi:10.1093/nar/gkp357 (2009).

29 Manoel-Caetano, F. S., Rossi, A. F. T., Calvet de Morais, G., Severino, F. E. & Silva, A. E. Upregulation of the APE1 and H2AX genes and miRNAs involved in DNA damage response and repair in gastric cancer. Genes Dis 6, 176–184, doi:10.1016/j.gendis.2019.03.007 (2019).

30 Yuan, C. L. et al. APE1 overexpression is associated with poor survival in patients with solid tumors: a meta-analysis. Oncotarget 8, 59720–59728, doi:10.18632/oncotarget.19814 (2017).

31 Wen, X. et al. APE1 overexpression promotes the progression of ovarian cancer and serves as a potential therapeutic target. Cancer Biomark 17, 313–322, doi:10.3233/CBM-160643 (2016).

32 Kumar, S. et al. Role of apurinic/apyrimidinic nucleases in the regulation of homologous recombination in myeloma: mechanisms and translational significance. Blood Cancer J 8, 92, doi:10.1038/s41408-018-0129-9 (2018).

33 Gafter-Gvili, A. et al. Oxidative stress-induced DNA damage and repair in human peripheral blood mononuclear cells: protective role of hemoglobin. PLoS One 8, e68341, doi:10.1371/journal.pone.0068341 (2013).

34 Helgason, H. et al. Loss-of-function variants in ATM confer risk of gastric cancer. Nat Genet 47, 906–910, doi:10.1038/ng.3342 (2015).

35 Li, X. et al. A comparison of per sample global scaling and per gene normalization methods for differential expression analysis of RNA-seq data. PLoS One 12, e0176185, doi:10.1371/journal.pone.0176185 (2017).

36 Ramos, M., Schiffer, L. & Waldron, L. TCGAutils: TCGA utility functions for data management. R package version 1.4.0. (2019).

37 Wickham, H. ggplot2: Elegant Graphics for Data Analysis. Springer-Verlag New York. (2016).

38 Xie, Y., Allaire, J. J. & Grolemund, G. R Markdown: The Definitive Guide. Chapman and Hall/CRC. (2018).

39 Allaire, J. J. et al. R: rmarkdown: Dynamic Documents for R. R package version 1.14. (2019).

40 RStudio Team. RStudio: Integrated Development for R. RStudio, Inc., Boston, MA. (2015).

41 R Core Team. R: A language and environment for statistical computing. R Foundation for Statistical Computing, Vienna, Austria. (2013).

42 Lin, Y. et al. APE1 senses DNA single-strand breaks for repair and signaling. Nucleic Acids Res, doi:10.1093/nar/gkz1175 (2019).

43 Cupello, S., Lin, Y. & Yan, S. Distinct roles of XRCC1 in genome integrity in Xenopus egg extracts. Biochem J 476, 3791–3804, doi:10.1042/BCJ20190798 (2019).

44 Hadi, M. Z. & Wilson, D. M., 3rd. Second human protein with homology to the Escherichia coli abasic endonuclease exonuclease III. Environ Mol Mutagen 36, 312–324, doi:http://dx.doi.org/10.1002/1098-2280(2000)36:4<312::AID-EM7>3.0.CO;2-K (2000).

45 Li, F. et al. Apn2 resolves blocked 3’ ends and suppresses Top1-induced mutagenesis at genomic rNMP sites. Nat Struct Mol Biol 26, 155–163, doi:10.1038/s41594-019-0186-1 (2019).

46 Johnson, R. E. et al. Identification of APN2, the Saccharomyces cerevisiae homolog of the major human AP endonuclease HAP1, and its role in the repair of abasic sites. Genes Dev. 12, 3137–3143, doi:10.1101/gad.12.19.3137 (1998).

47 Unk, I. et al. Stimulation of 3’-->5’ exonuclease and 3’-phosphodiesterase activities of yeast apn2 by proliferating cell nuclear antigen. Mol Cell Biol 22, 6480–6486, doi:10.1128/mcb.22.18.6480-6486.2002 (2002).

48 Ribar, B., Izumi, T. & Mitra, S. The major role of human AP-endonuclease homolog Apn2 in repair of abasic sites in Schizosaccharomyces pombe. Nucleic Acids Res 32, 115–126, doi:10.1093/nar/gkh151 (2004).

49 Ma, W., Resnick, M. A. & Gordenin, D. A. Apn1 and Apn2 endonucleases prevent accumulation of repair-associated DNA breaks in budding yeast as revealed by direct chromosomal analysis. Nucleic Acids Res 36, 1836–1846, doi:10.1093/nar/gkm1148 (2008).

50 Tsuchimoto, D. et al. Human APE2 protein is mostly localized in the nuclei and to some extent in the mitochondria, while nuclear APE2 is partly associated with proliferating cell nuclear antigen. Nucleic Acids Res 29, 2349–2360, doi:10.1093/nar/29.11.2349 (2001).

51 Burkovics, P., Szukacsov, V., Unk, I. & Haracska, L. Human Ape2 protein has a 3’-5’ exonuclease activity that acts preferentially on mismatched base pairs. Nucleic Acids Res 34, 2508–2515, doi:10.1093/nar/gkl259 (2006).

52 Mengwasser, K. E. et al. Genetic Screens Reveal FEN1 and APEX2 as BRCA2 Synthetic Lethal Targets. Mol Cell 73, 885–899 e886, doi:10.1016/j.molcel.2018.12.008 (2019).

53 Ide, Y. et al. Growth retardation and dyslymphopoiesis accompanied by G2/M arrest in APEX2-null mice. Blood 104, 4097–4103, doi:10.1182/blood-2004-04-1476 (2004).

54 Guikema, J. E. et al. APE1- and APE2-dependent DNA breaks in immunoglobulin class switch recombination. J Exp Med 204, 3017–3026, doi:10.1084/jem.20071289 (2007).

55 Guikema, J. E. et al. Apurinic/apyrimidinic endonuclease 2 is necessary for normal B cell development and recovery of lymphoid progenitors after chemotherapeutic challenge. J Immunol 186, 1943–1950, doi:10.4049/jimmunol.1002422 (2011).

56 Permata, T. B. M. et al. Base excision repair regulates PD-L1 expression in cancer cells. Oncogene 38, 4452–4466, doi:10.1038/s41388-019-0733-6 (2019).

57 Sun, L. L. et al. Inhibition of ATR downregulates PD-L1 and sensitizes tumor cells to T cell-mediated killing. Am J Cancer Res 8, 1307–1316 (2018).

58 Vendetti, F. P. et al. ATR kinase inhibitor AZD6738 potentiates CD8+ T cell-dependent antitumor activity following radiation. J Clin Invest 128, 3926–3940, doi:10.1172/JCI96519 (2018).

59 Slyskova, J. et al. Functional, genetic, and epigenetic aspects of base and nucleotide excision repair in colorectal carcinomas. Clin Cancer Res 18, 5878–5887, doi:10.1158/1078-0432.CCR-12-1380 (2012).

60 Vodenkova, S. et al. Base excision repair capacity as a determinant of prognosis and therapy response in colon cancer patients. DNA Repair (Amst) 72, 77–85, doi:10.1016/j.dnarep.2018.09.006 (2018).

