## Supplemental Figures for "Genomic alterations and abnormal expression of APE2 in multiple cancers"

**Supplementary Figures and Legend:**

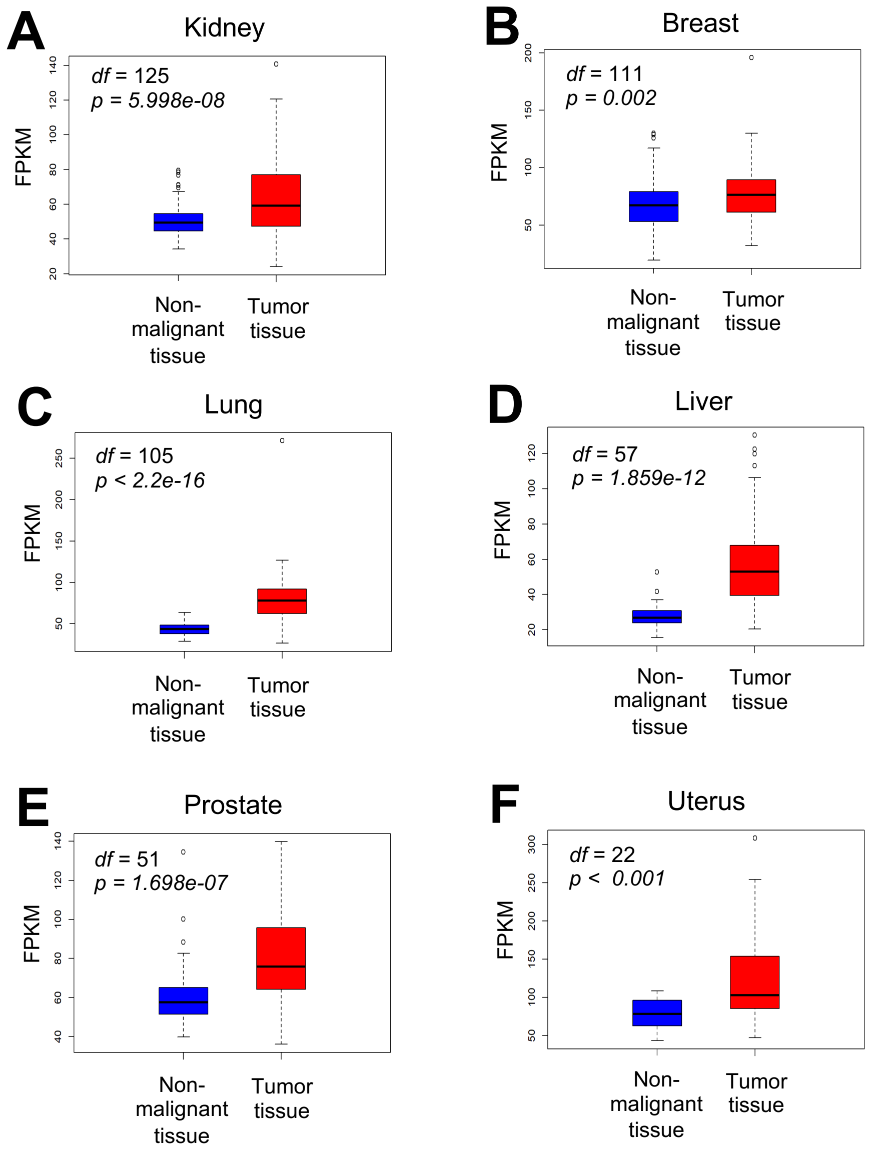


**Fig. S1**. APE1 mRNA expression between tumor tissue and matched non-malignant tissue per individual from 6 different cancer types including kidney (**A**), breast (**B**), lung (**C**), liver (**D**), prostate (**E**), and uterus (**F**).

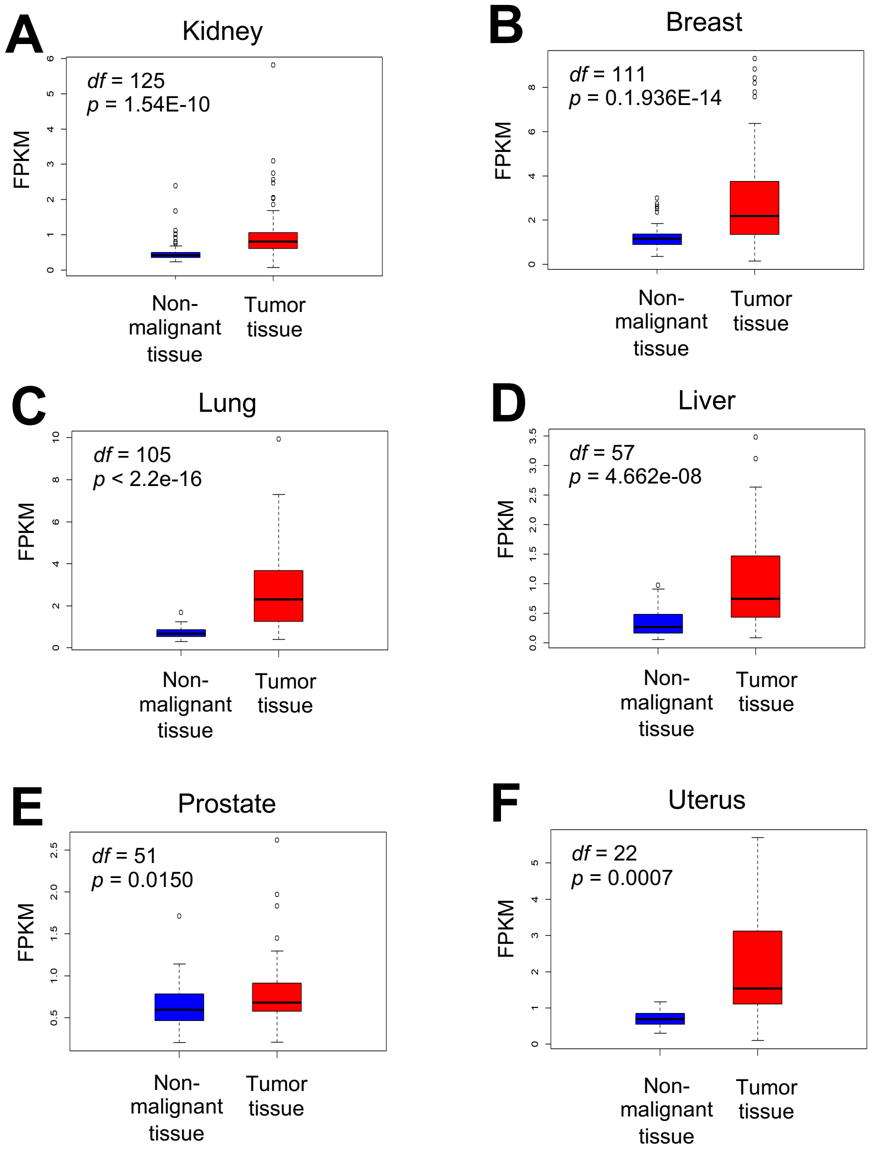


**Fig. S2**. BRCA1 mRNA expression between tumor tissue and matched non-malignant tissue per individual from 6 different cancer types including kidney (**A**), breast (**B**), lung (**C**), liver (**D**), prostate (**E**), and uterus (**F**).

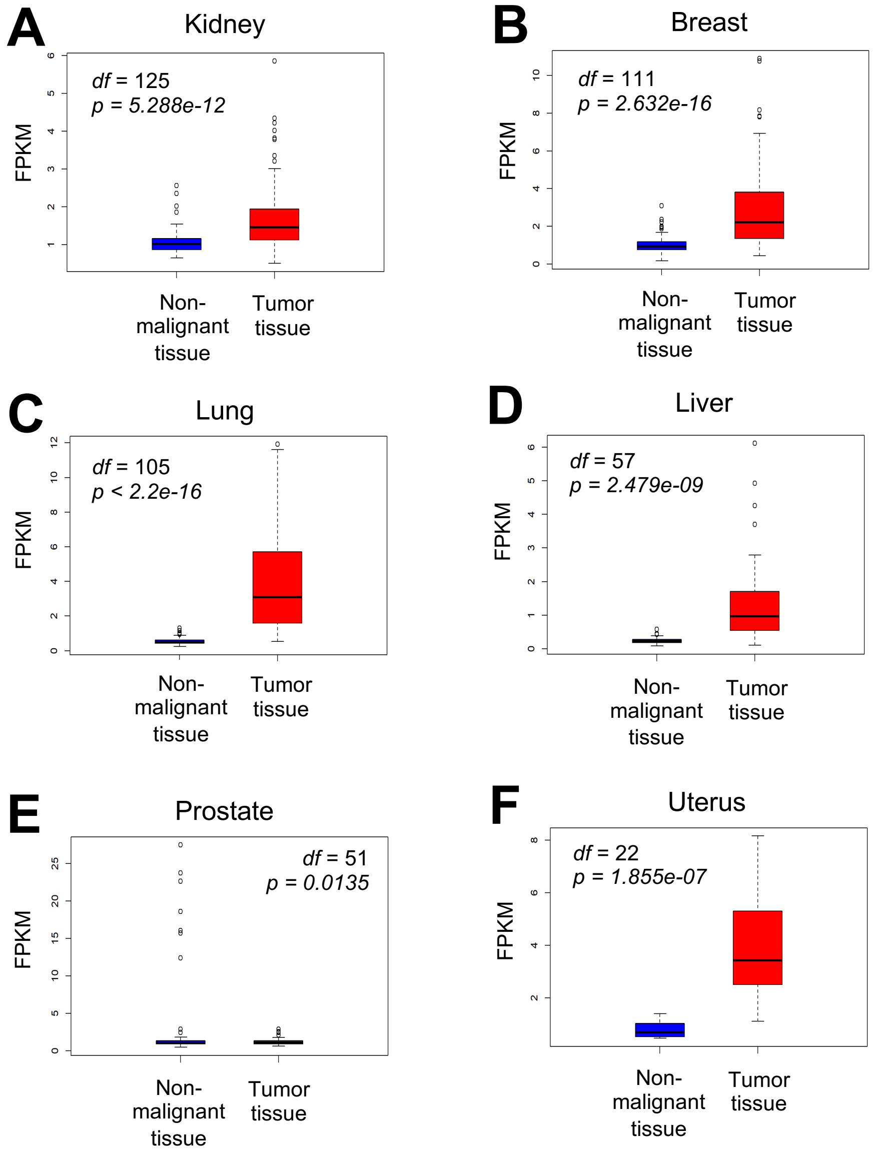


**Fig. S3**. Chk1 mRNA expression between tumor tissue and matched non-malignant tissue per individual from 6 different cancer types including kidney (**A**), breast (**B**), lung (**C**), liver (**D**), prostate (**E**), and uterus (**F**).

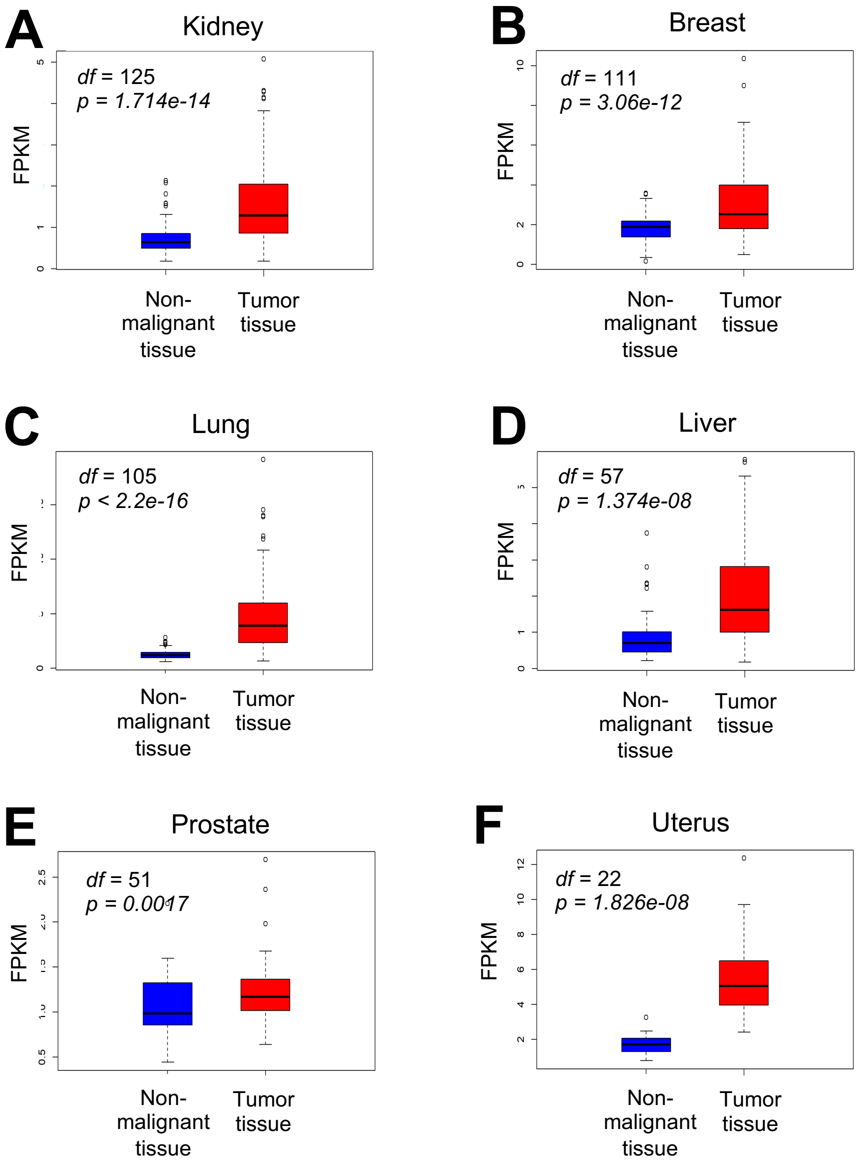


**Fig. S4**. Chk2 mRNA expression between tumor tissue and matched non-malignant tissue per individual from 6 different cancer types including kidney (**A**), breast (**B**), lung (**C**), liver (**D**), prostate (**E**), and uterus (**F**).

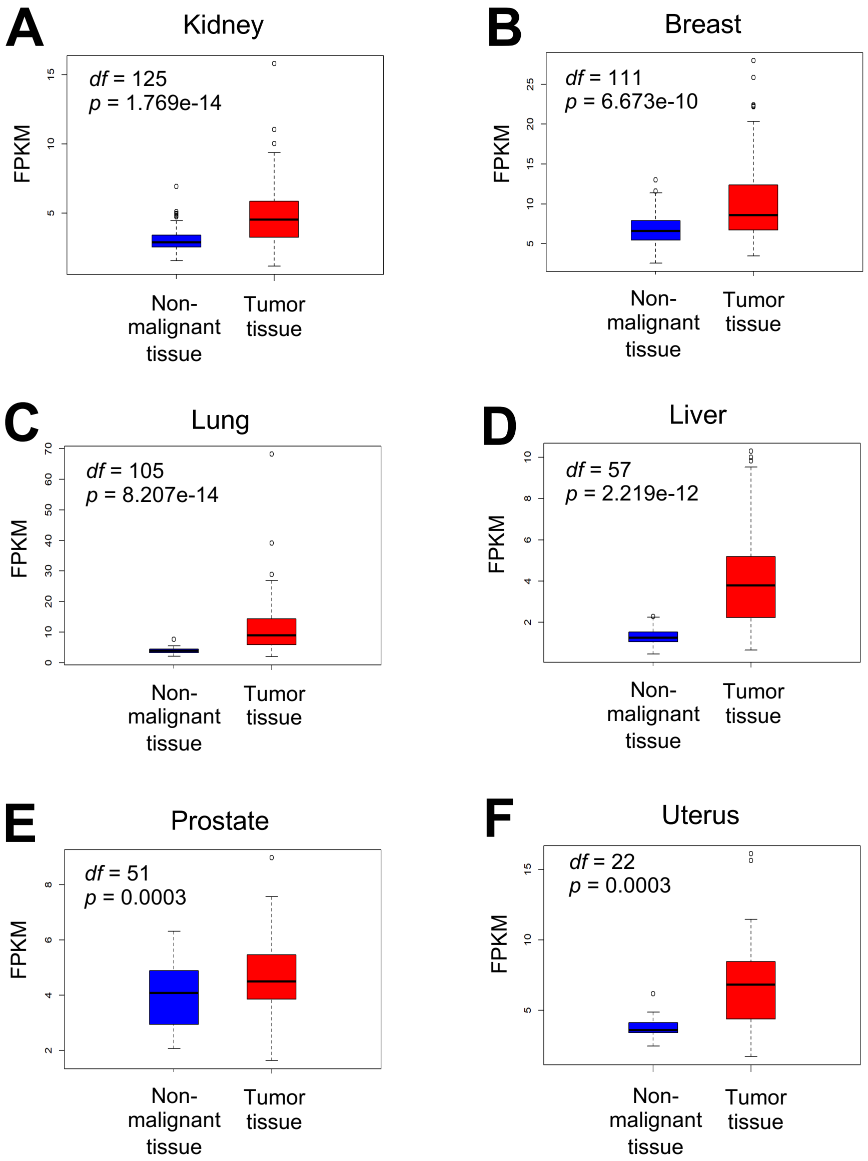


**Fig. S5**. TopBP1 mRNA expression between tumor tissue and matched non-malignant tissue per individual from 6 different cancer types including kidney (**A**), breast (**B**), lung (**C**), liver (**D**), prostate (**E**), and uterus (**F**).

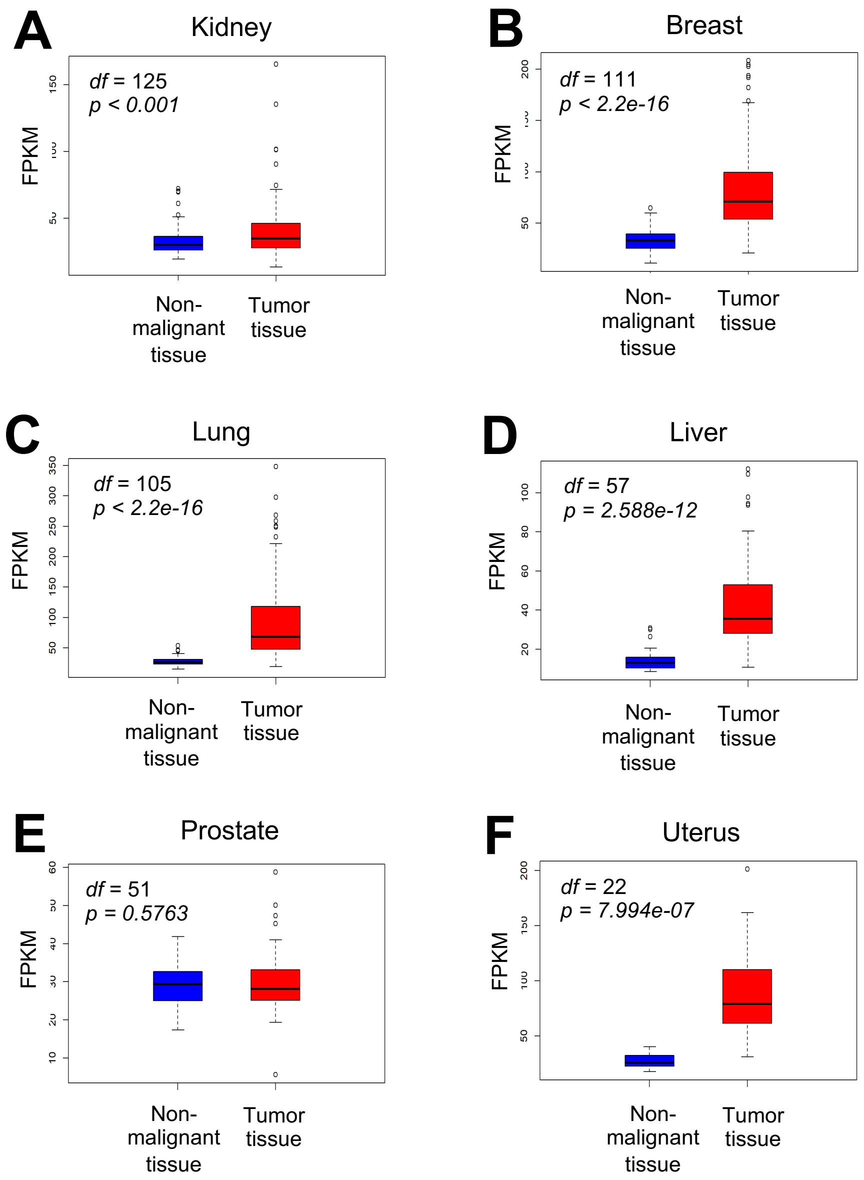


**Fig. S6**. PCNA mRNA expression between tumor tissue and matched non-malignant tissue per individual from 6 different cancer types including kidney (**A**), breast (**B**), lung (**C**), liver (**D**), prostate (**E**), and uterus (**F**).

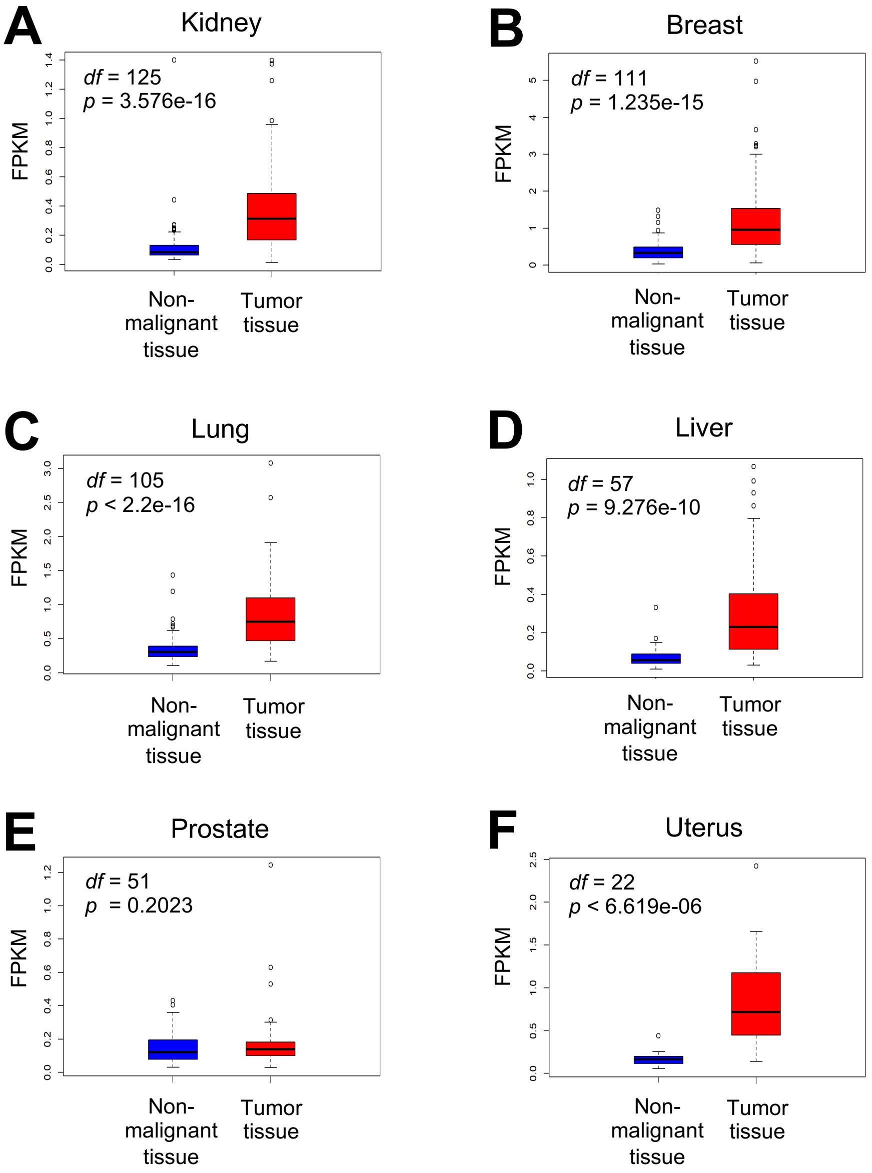


**Fig. S7**. BRCA2 mRNA expression between tumor tissue and matched non-malignant tissue per individual from 6 different cancer types including kidney (**A**), breast (**B**), lung (**C**), liver (**D**), prostate (**E**), and uterus (**F**).

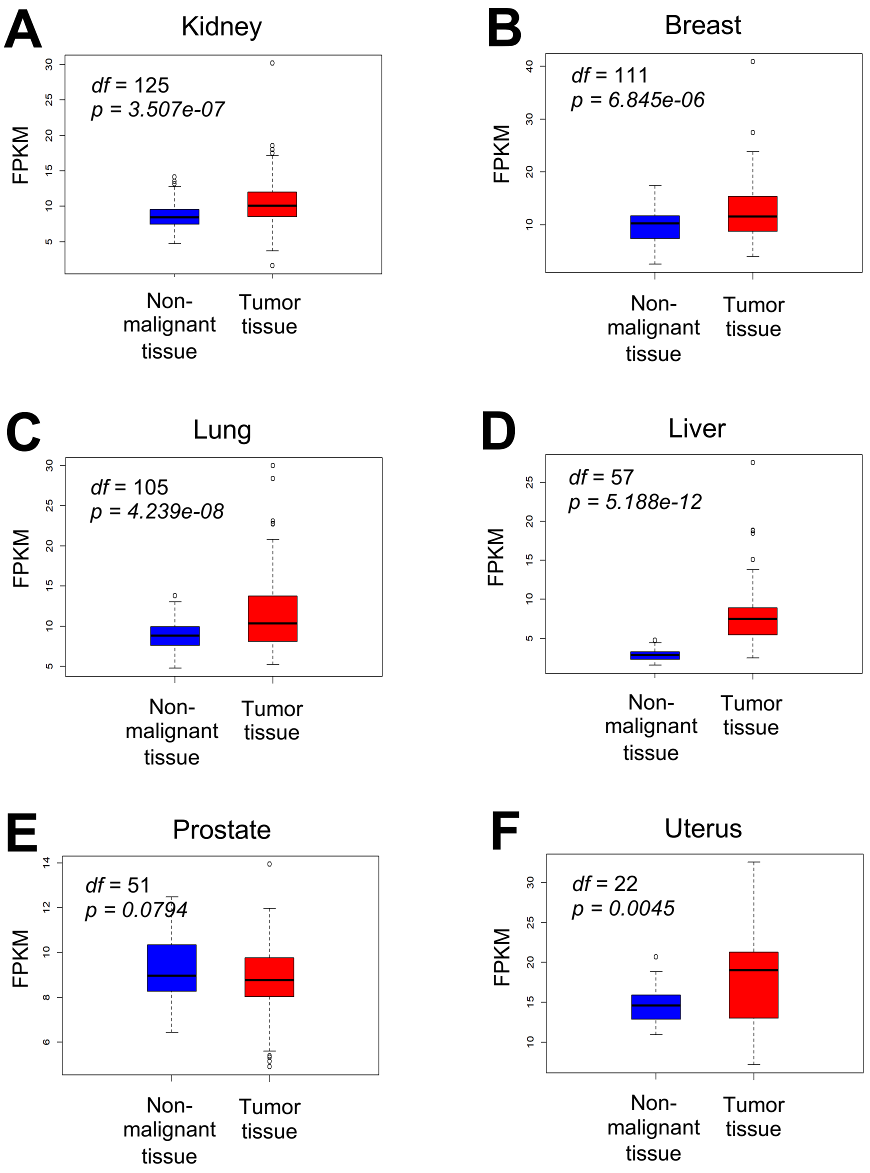


**Fig. S8.** XRCC1 mRNA expression between tumor tissue and matched non-malignant tissue per individual from 6 different cancer types including kidney (**A**), breast (**B**), lung (**C**), liver (**D**), prostate (**E**), and uterus (**F**).

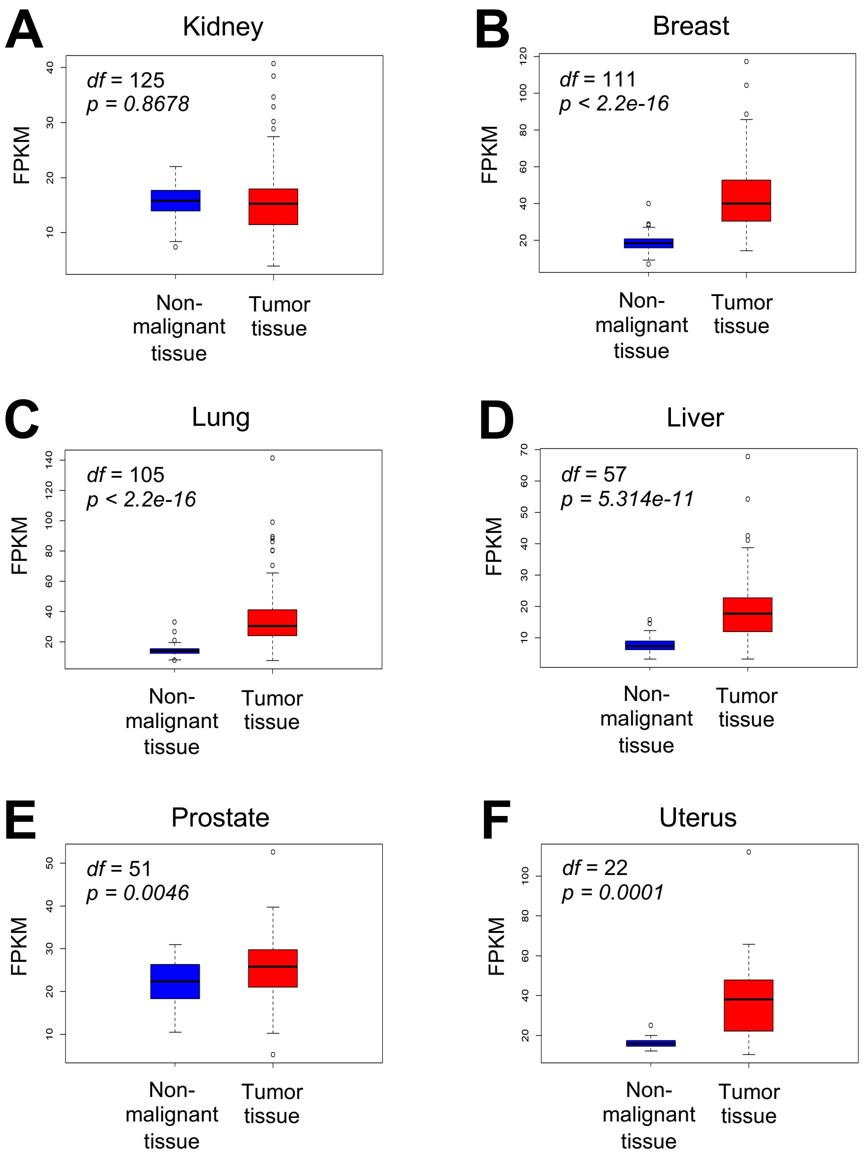


**Fig. S9**. PARP1 mRNA expression between tumor tissue and matched non-malignant tissue per individual from 6 different cancer types including kidney (**A**), breast (**B**), lung (**C**), liver (**D**), prostate (**E**), and uterus (**F**).

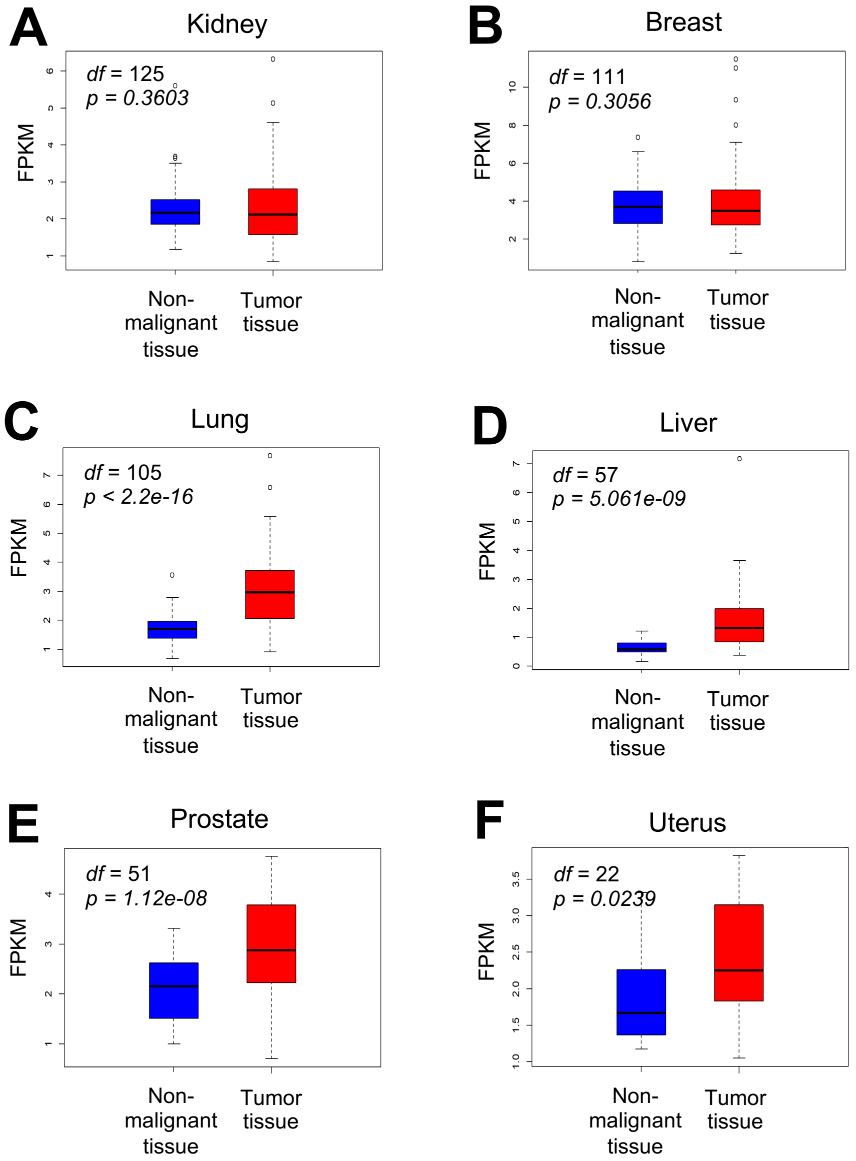


**Fig. S10**. ATR mRNA expression between tumor tissue and matched non-malignant tissue per individual from 6 different cancer types including kidney (**A**), breast (**B**), lung (**C**), liver (**D**), prostate (**E**), and uterus (**F**).

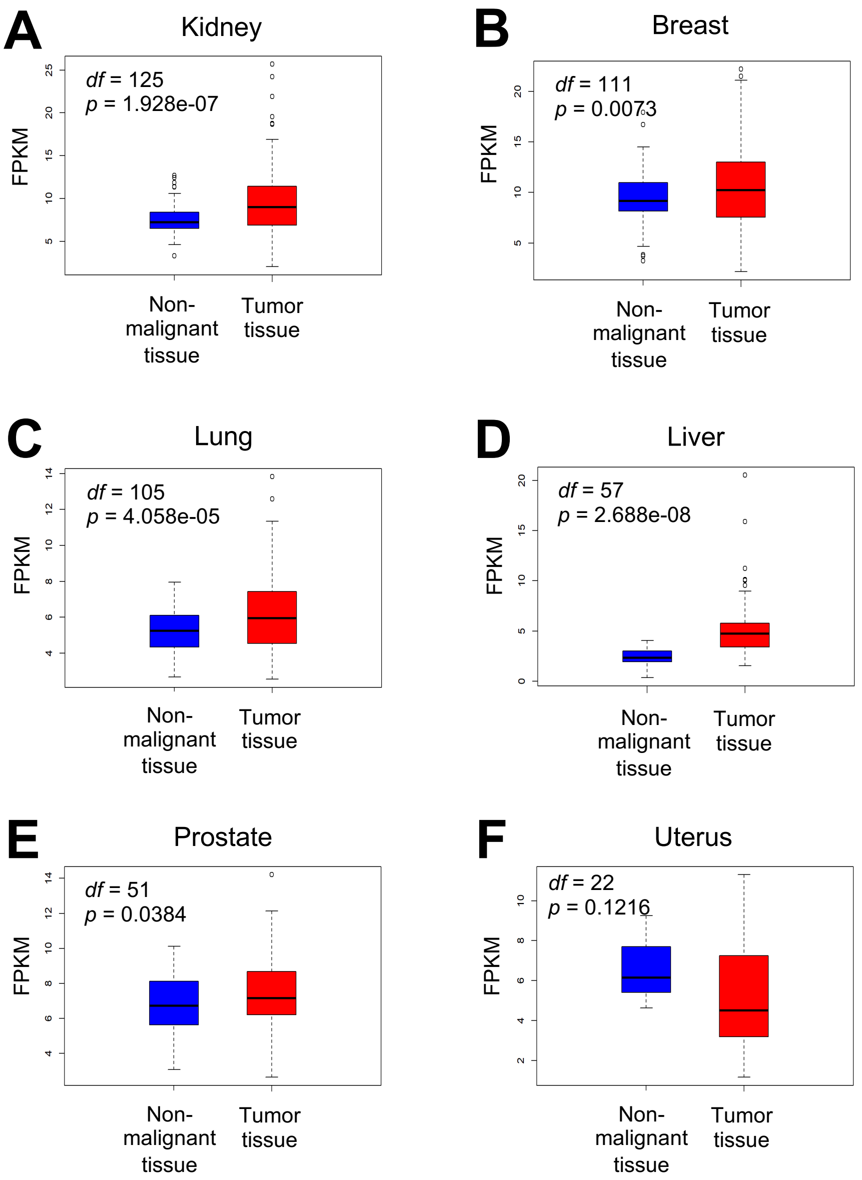


**Fig. S11**. Rad50 mRNA expression between tumor tissue and matched non-malignant tissue per individual from 6 different cancer types including kidney (**A**), breast (**B**), lung (**C**), liver (**D**), prostate (**E**), and uterus (**F**).

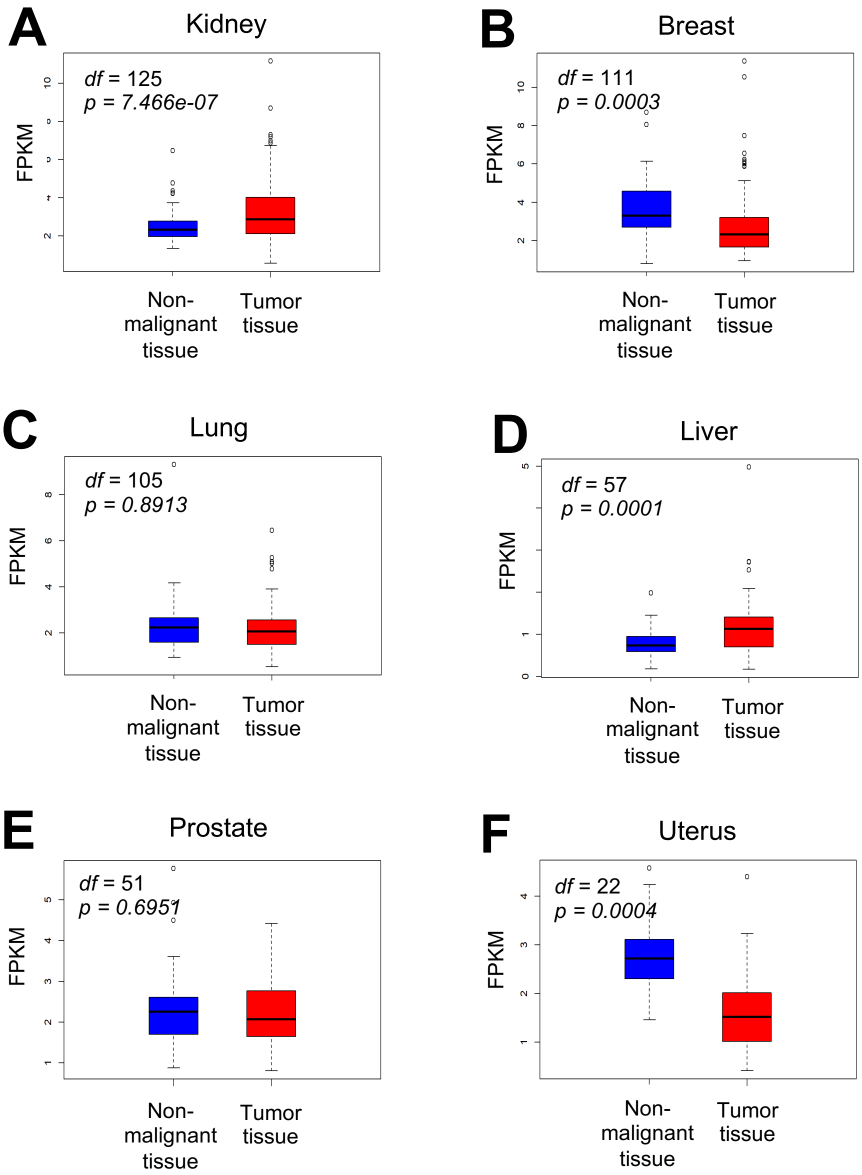


**Fig. S12**. ATM mRNA expression between tumor tissue and matched non-malignant tissue per individual from 6 different cancer types including kidney (**A**), breast (**B**), lung (**C**), liver (**D**), prostate (**E**), and uterus (**F**).

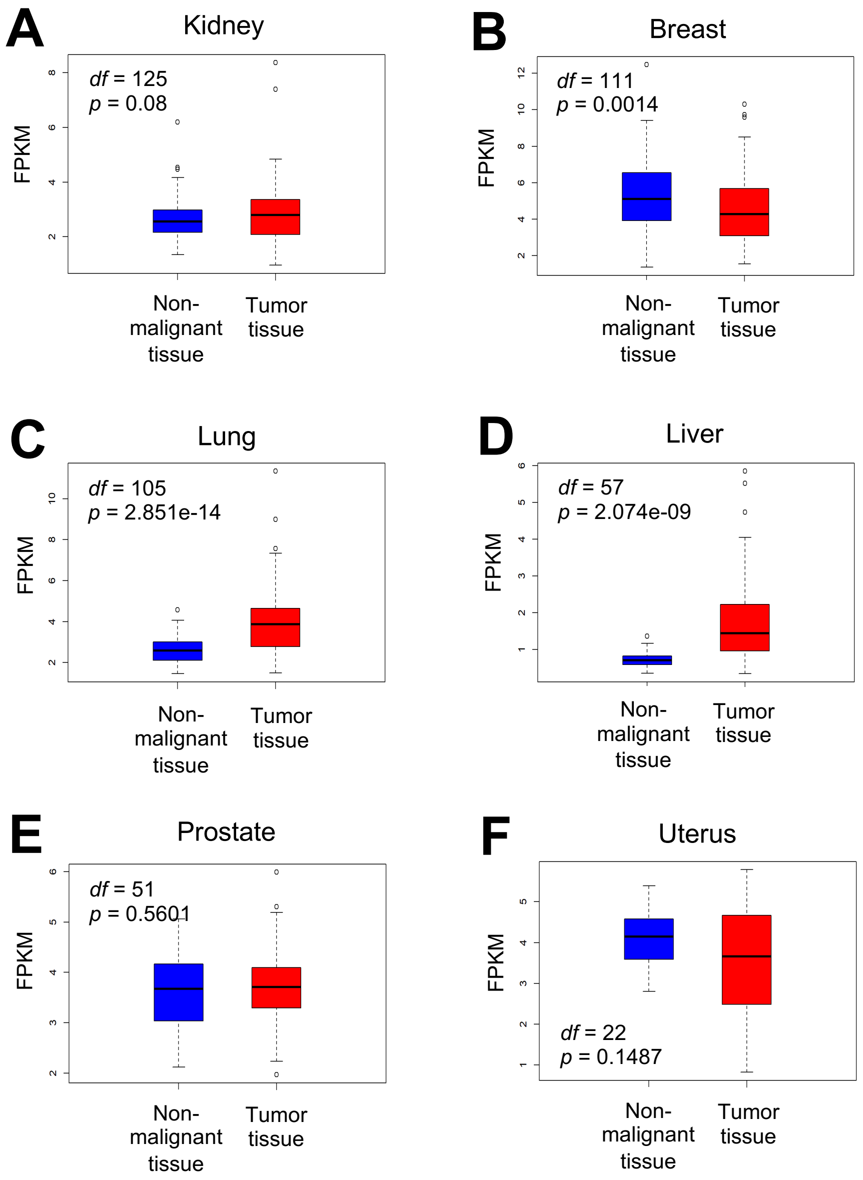


**Fig. S13**. Mre11 mRNA expression between tumor tissue and matched non-malignant tissue per individual from 6 different cancer types including kidney (**A**), breast (**B**), lung (**C**), liver (**D**), prostate (**E**), and uterus (**F**).

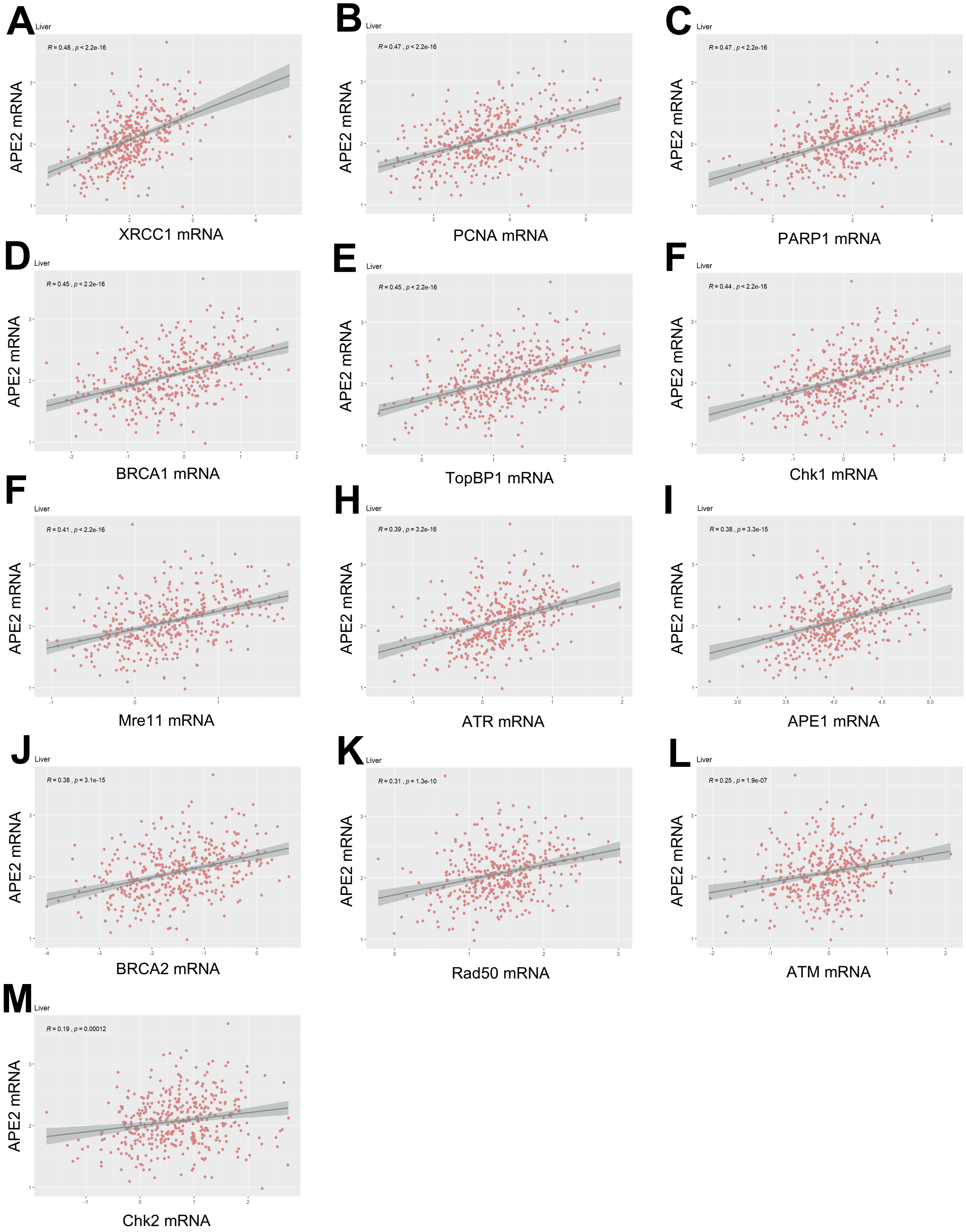


**Fig. S14.** Correlation between mRNA expression of APE2 and 13 other DNA repair and DDR proteins in tumor tissues of liver cancer. *R* and *p* values are listed in each panel.

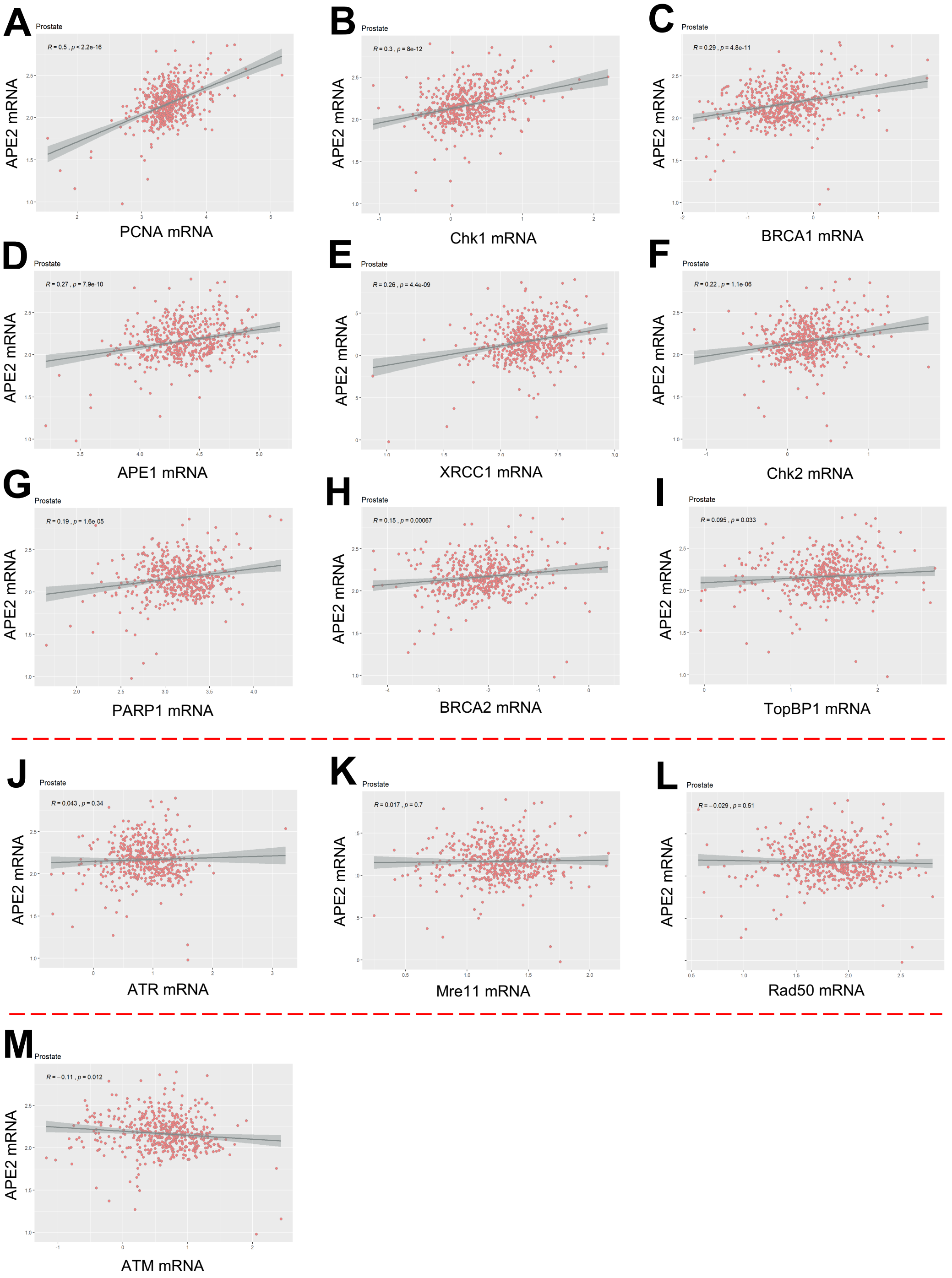


**Fig. S15.** Correlation between mRNA expression of APE2 and 13 other DNA repair and DDR proteins in tumor tissues of prostate cancer. *R* and *p* values are listed in each panel.

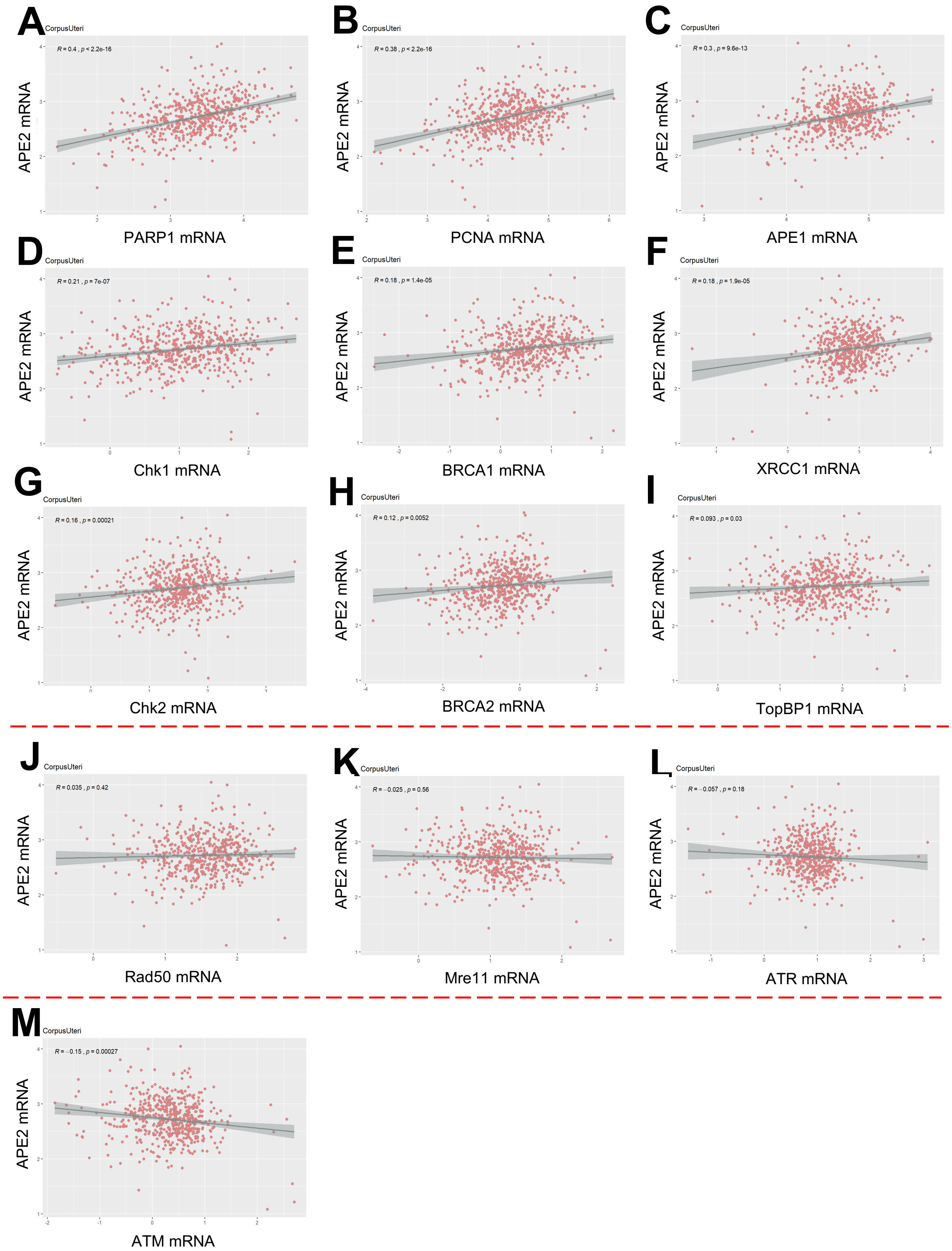


**Fig. S16.** Correlation between mRNA expression of APE2 and 13 other DNA repair and DDR proteins in tumor tissues of uterine cancer. *R* and *p* values are listed in each panel.

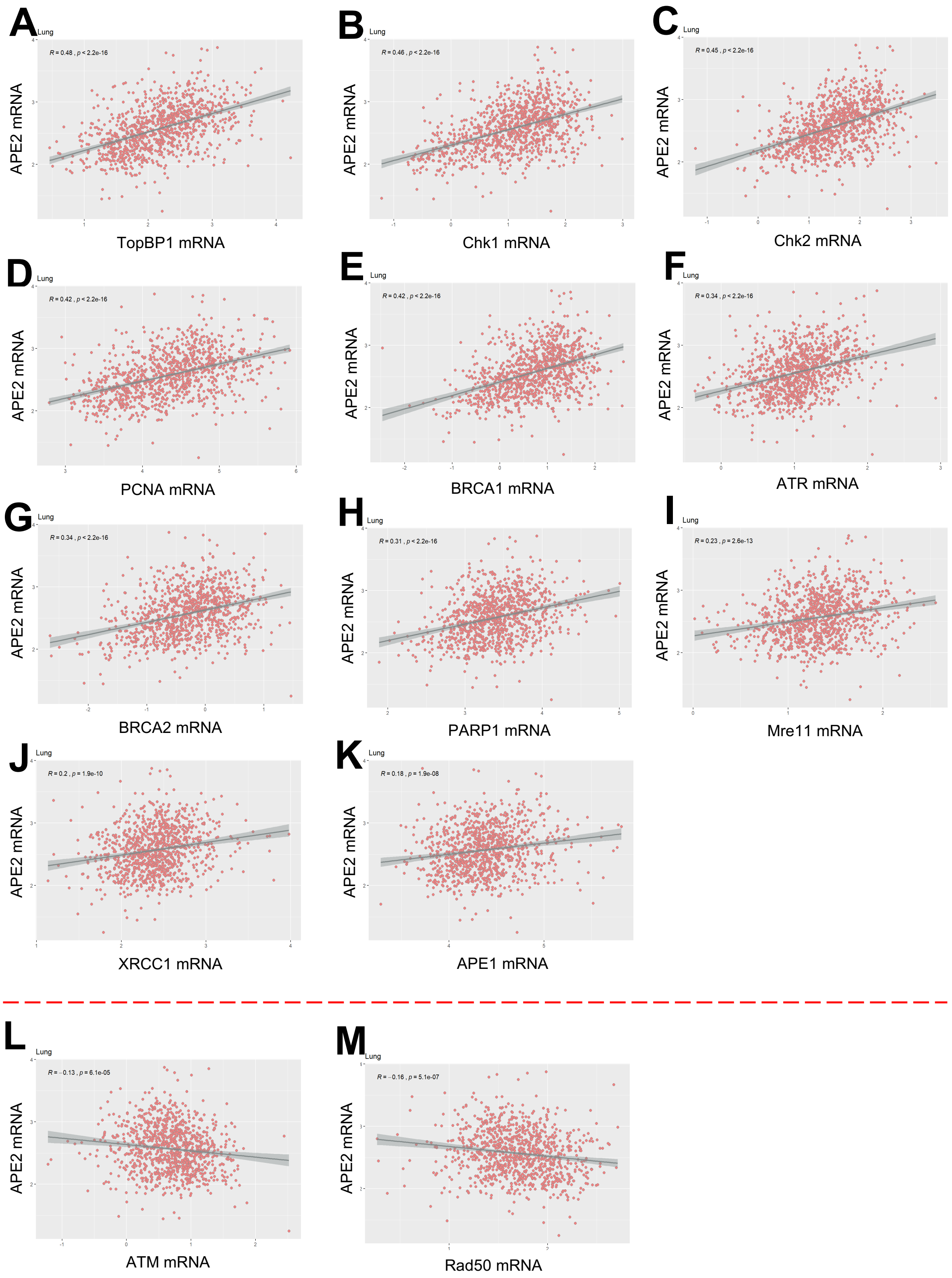


**Fig. S17.** Correlation between mRNA expression of APE2 and 13 other DNA repair and DDR proteins in tumor tissues of lung cancer. *R* and *p* values are listed in each panel.

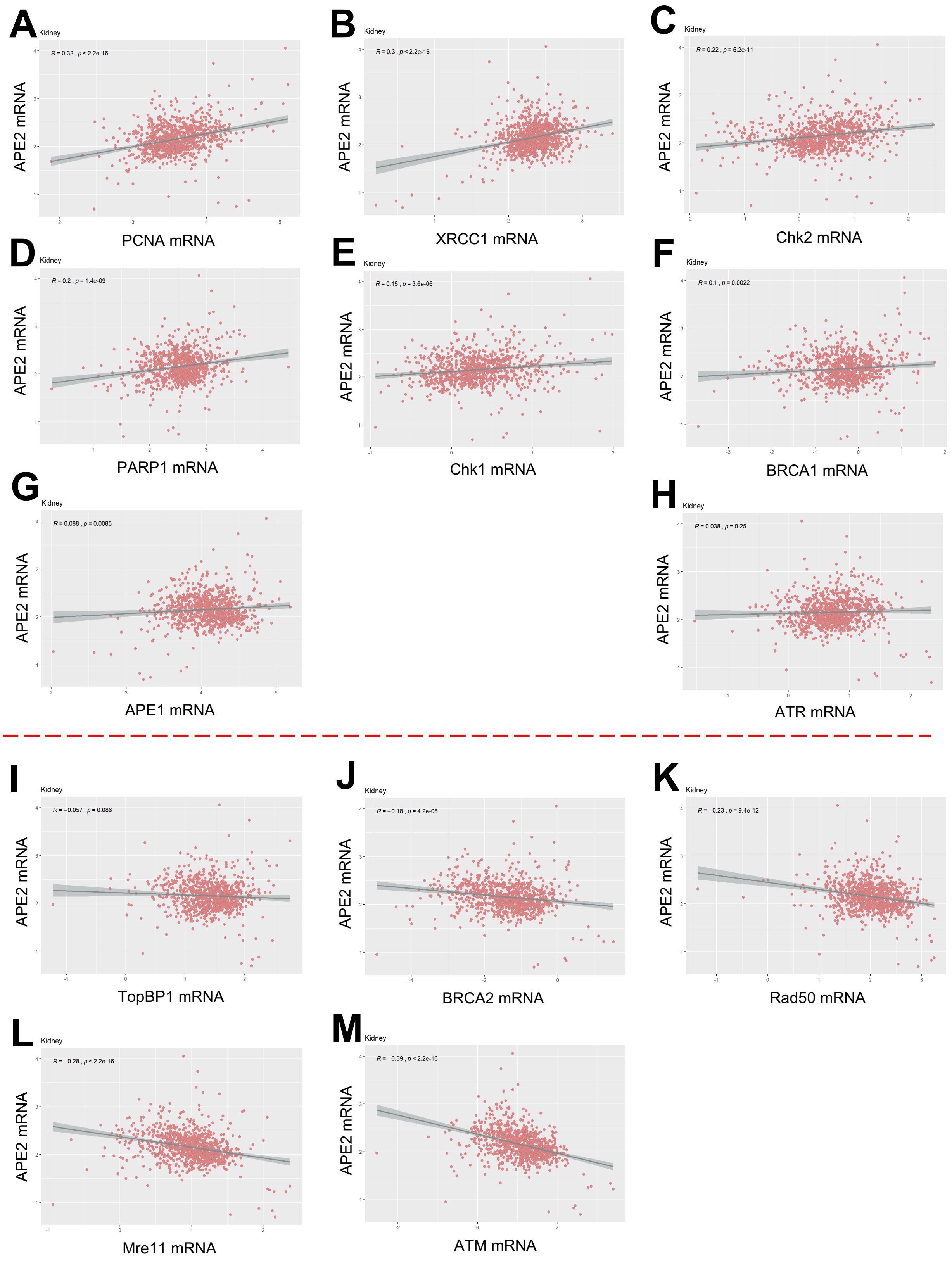


**Fig. S18.** Correlation between mRNA expression of APE2 and 13 other DNA repair and DDR proteins in tumor tissues of kidney cancer. *R* and *p* values are listed in each panel.
